# Canonical host-pathogen tradeoffs subverted by mutations with dual benefits

**DOI:** 10.1101/818492

**Authors:** Robert Beardmore, Mark Hewlett, Rafael Peña-Miller, Carlos Reding, Ivana Gudelj, Justin R. Meyer

## Abstract

Tradeoffs between life history traits impact diverse biological phenomena, including the maintenance of biodiversity. We sought to study two canonical tradeoffs in a model host-parasite system consisting of bacteriophage lambda and *Escherichia coli*: i) parasite resistance for growth and ii) phage infectivity for host-range. We report that these previously hypothesised tradeoffs are, in fact, tradeups. While the observation of tradeups was surprising, they should be expected because if traits X and Y tradeoff, so too traits Y and Z, then X and Z will tradeup. By considering five different *E. coli* trait correlations we uncovered several tradeups and tradeoffs. Using mathematical models, we establish that tradeups need not inhibit biodiversity, as previously thought, and can help maintain it through high-dimensional trait interactions. We provide a mechanistic explanation for how tradeups emerge and give reasons for why tradeups can even evolve in well-adapted genomes.

All data will be posted at https://github.com/rebear217 and mirrored at http://people.exeter.ac.uk/reb217/rebHomePage/data.html on acceptance.

## Introduction

Life history tradeoffs and parasitism are key to our understanding of biodiversity. Different theoretical frameworks^1–4^ predict, quite consistently, that if improvements in one trait come with costs in a second trait and if the geometry of that cost has the right form, polymorphisms are maintained. At its logical limit, this suggests tradeoff geometries might be inferred from biodiversity patterns.^4^ Infection by parasites is a key driver of genetic diversity and one parasitism tradeoff is central to our understanding of how parasites target their hosts: the host-range tradeoff (HRTO). The HRTO postulates that if a parasite is efficient at targeting one host, it will be inefficient at targeting others. As a result, a lock and key interaction should describe host-parasite interactions^5–7^ whereby each parasitic ‘key’ can only ‘unlock’ a subset of hosts.

Phage therapy represents an important potential adjuvant to antibiotic therapy for life-threatening diseases^8–10^ and tradeoffs are important for the future success of phage therapy. For if a pathogenic bacterium evolves that is able to evade all the phage that it is currently susceptible to, that bacterium will no longer be treatable by phage. However, a postulated cost of resistance tradeoff^11^ (CORT) whereby increases in phage resistance come only with reductions in bacterial growth rate suggests that such highly-resistant bacteria, which are observed in patients,^9^ will be poor replicators.

Given the large number of postulated tradeoffs in the literature^12–20^ and their importance in ecology and medicine, we used an *in vitro* coevolutionary model system to test for the presence of (1) the parasite’s HRTO and (2) a bacterial CORT. We also tested for (3) the rate-yield tradeoff^21^ (RYTO) whereby bacteria that replicate more quickly do so less efficiently.^22^

For this we implemented a host-parasite model system of *Escherichia coli* and phage *λ* to create a library of phage resistant hosts and parasites that vary in their host range. Isogenic cultures of each were co-cultured and allowed to coevolve for three days, just enough time for the bacteria to evolve resistance and for the phage to counter, thus 47 distinct bacterial host genotypes that vary by mutations in the viral receptor (maltoporin LamB) and 93 viral genotypes were isolated (see Methods). We observed a within-strain RYTO for almost all bacterial strains, consistent with prior expectations.^3^ But our main finding is this: the library of over 4,000 host-parasite interactions exhibited little tradeoff data. There is no between-strain RYTO, no CORT, and a tradeup appeared instead of the HRTO.

Unexpectedly, the host range data from the library took the form a trade-up whereby phage that are more infectious to any given host can also infect more hosts. We therefore observed no lock and key as some viruses were simply more infectious than others. Similarly, bacteria evolved high levels of resistance and paid no discernible fitness cost for their resistance. We even observed extreme cases whereby certain bacteria were pan-resistant: they could resist attack from all *λ* genotypes in the library and yet they grew more quickly than the phage-susceptible wild type from which they were derived. Thus some evolved bacteria have dual adaptive benefits relative to their wild-type ancestor.

One interpretation of this result could be that the ancestral genome is not operating at its limits in terms of replication and resistance where the tradeoffs would be observed. Further investigations into the mechanisms supporting these dual benefits revealed an alternative explanation. By analysing structural changes in the phage receptor on the bacterial cell surface, LamB, and by analysing tradeoffs mediated by these changes, we discovered a relationship between phage resistance, nutrient uptake rate, growth rate and yield that can produce a tradeup between resistance and absolute fitness. Using theory and data, we therefore argue that multiple, interacting tradeoffs can conspire to bring about dual benefits (Supplementary §6) and, as a result, tradeups could be just as likely in polymorphic biological systems as tradeoffs.

## Results

Tradeoffs and parasitism are central to biodiversity theory because host-parasite coevolution combined with tradeoffs can produce striking levels of biodiversity. Figure 1(a) illustrates this using a theoretical bacteria-phage model. Here, simulations of a lock and key infection mechanism produced Red Queen dynamics where the two species engage in a game of hide-and-seek within the space of possible bacterial and phage phenotypes: phage evolve to seek new bacterial hosts and bacteria coevolve to hide from contemporary phage. Bacterial mutations with just small enough phenotypic effect to disappear from view of those phage, say by varying a cell surface receptor, allow the mutant host to avoid infection and so persist, even if it pays a fitness cost to achieve this. Phage then seek out these hidden hosts through small effect mutations of their own, creating polymorphic populations of specialist hosts and parasites where each of the latter provides a niche for the former, at least, that is, until more mutants arise.

**Figure 1:**
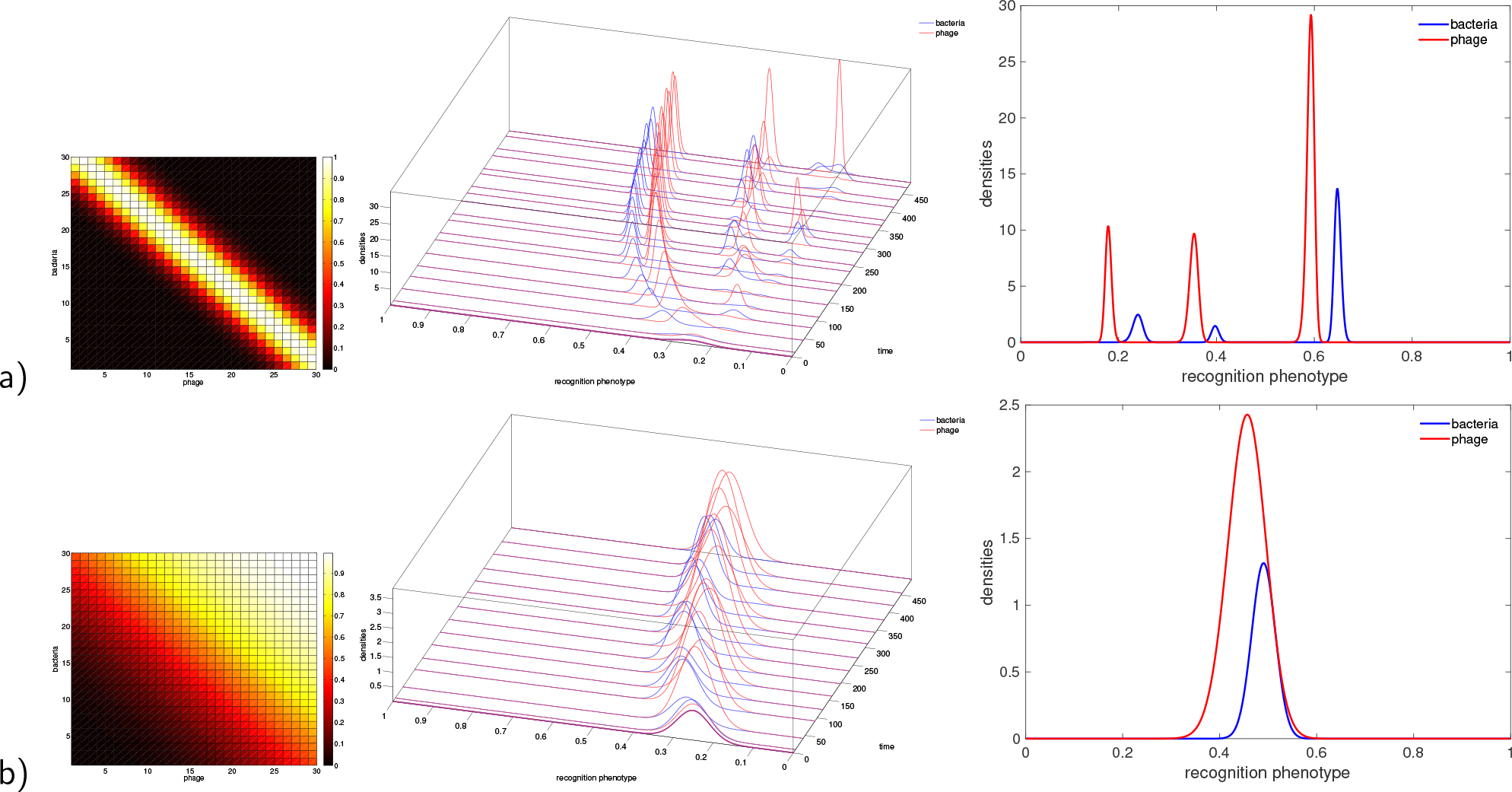
A bacteria-phage coevolutionary model (4) simulated under two different assumptions (a) a lock and key infection matrix and (b) a graded matrix: the former generates a multiplicity of phage niches, the latter does not. a) The heat map (left column) illustrates the lock and key whereby each phage can infect a limited subset of all hosts, the centre plot shows the ensuring Red Queen dynamics, note the diversity it generates through phenotypic branching processes that arise from an initially monomorphic population, the final temporal state of which is shown in the rightmost plot. The latter shows the density of hosts and parasites at the last simulated timepoint where 3 phenotypic clusters of ‘matched’ bacteria and phage are visible. b) Analogous simulations to a) were performed, except now the graded infection matrix produces a Red Queen dynamic without diversification: phage and bacteria are monomorphic populations at all times. (See Methods for model details.)

Using this strain library we tested whether the *E. coli* and phage *λ* infection mechanism exhibit a lock and key structure^7^ (a.k.a. ‘modular’,^6, 23^ see [23, Figure 3]) and whether it possessed a HRTO whereby more specialist phage infect fewer hosts. Now, given two libraries of *N* bacteria and *M* phage, an *infection matrix* is an *N* × *M* array of likelihoods that bacterial genotype *i* can be infected by phage genotype *j* on contact.^23^ To obtain numerical proxies for these likelihoods (with phenotypes called *infectivities* for phage and *susceptibilities* for bacteria) we performed *N* infection assays (see Methods) on bacterial lawns, one lawn per bacterium inoculated with all *M* phage. Figure S1 details how the infectivities and susceptibilities are quantified using imaging algorithms.

**Figure 2:**
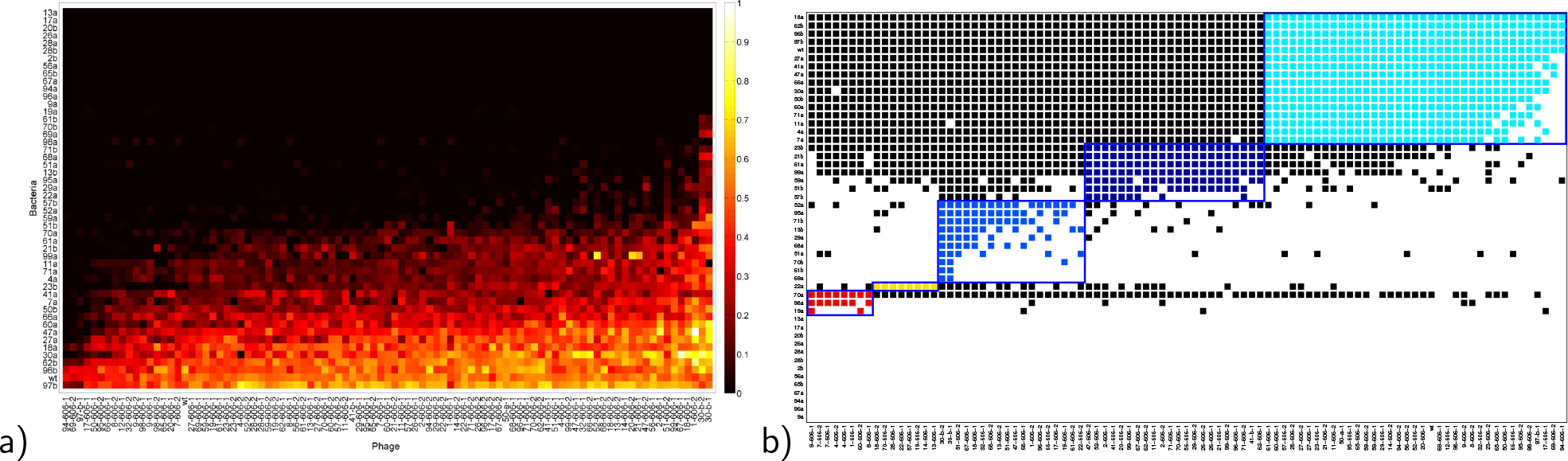
A bacteria-phage *λ* infection matrix: it is nested (i.e. graded), not modular (i.e. lock and key). a) A matrix of bacteria-phage interactions is shown as a heatmap where yellow and white are the most infectious phage-bacteria interactions whereas black show phage unable to infect the corresponding bacterium. The black region therefore represents a cluster of ‘pan-phage-resistant’ bacterial genotypes, namely bacteria that can be infected by none of the isolated phage. b) Attempts to categorise the matrix from (a) into modular sub-interactions using BiMat (and therefore determining its ‘lock and key’ sub-structures) fail as the matrix is highly nested (modularity metric *Q* ≈ 0.085 is no higher (indeed lower) than expected for random networks (rn) of the same dimension: mean(*Q*_*rn*_) ≈ 0.102, sd(*Q*_*rn*_) ≈ 0.01, *Z* ≈ −1.70, *n* = 1, 000 rns tested; a nestedness metric *N* ≈ 0.78 is significantly greater than expected for random networks: mean(*N*_*rn*_) ≈ 0.46, sd(*N*_*rn*_) ≈ 0.006, *Z* ≈ 58.6, *n* = 1, 000). This representation shows 5 optimised and coloured sub-structures on top of a binary (BW) colour scheme for pairwise interactions where black is used for non-zero interactions and white for zero; so the blocked colours indicate putative most-likely bacteria-phage interacting sub-modules. Note the black interactions are not contained within the coloured modules, highlighting an absence of locks and keys.

**Figure 3:**
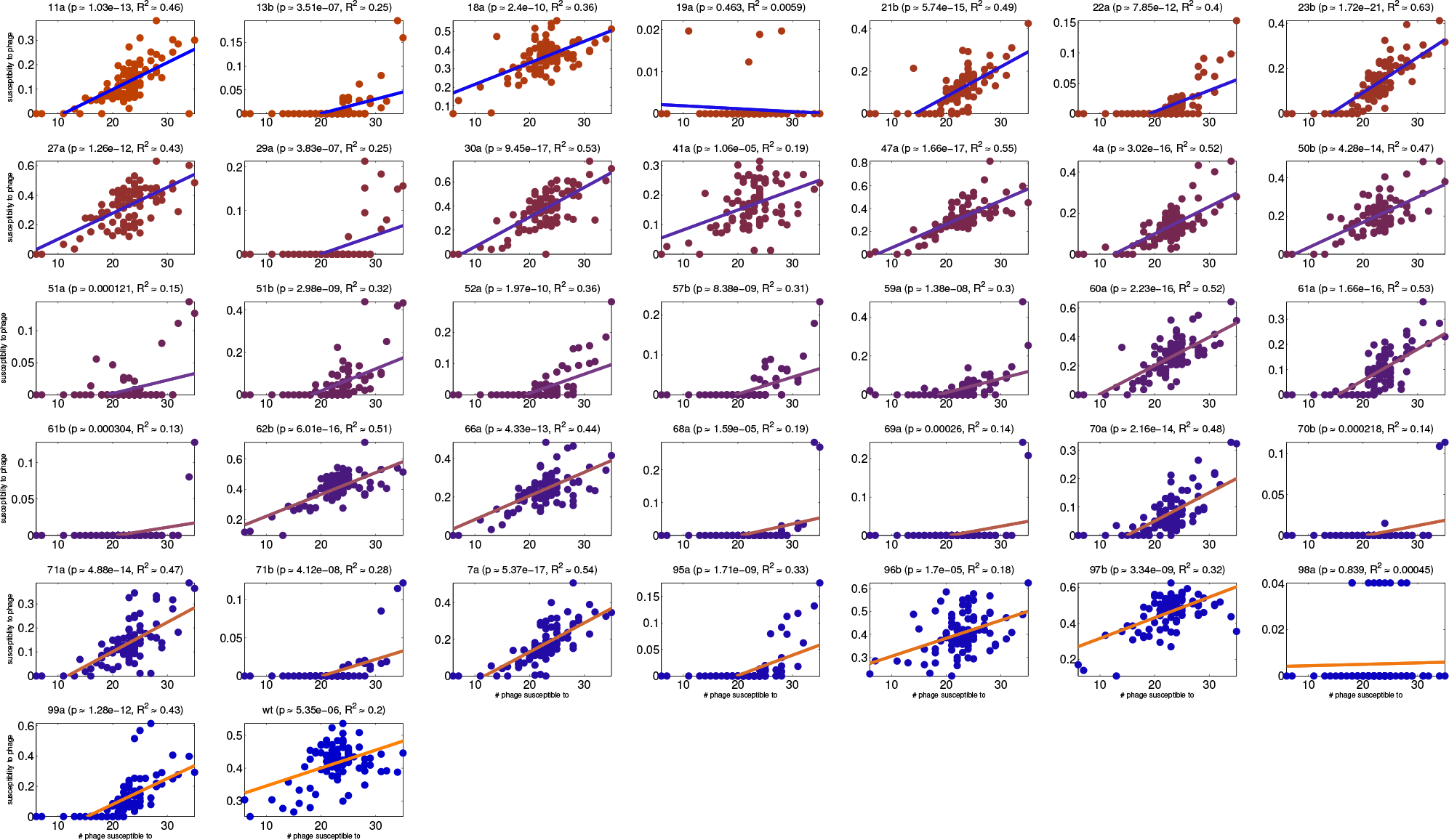
An infectivity-resistance tradeup: the smaller the set of phage that can infect an *E. coli* genotype, the less susceptible that genotype is to the phage that can infect it. In other words, the more susceptible an *E. coli* genotype is to each phage, the more phage there are that can infect it. We deduce this by performing linear regressions between the infectivity values in the imaging-derived matrix in Figure 2(a) and the number of infectious phage for each *E.coli* genotype: linear regressions for all phage genotypes are shown, the significance of the regression is given in the plot titles (all but two have significantly positive slopes).

Figure 2(a) shows the resulting *N* × *M* matrix, denoted Φ, of infectivities and sensitivities for the full library of phage and bacteria, displayed as a heat map. It provides no evidence of a lock and key structure where Φ would be ‘diagonally dominant’, like Figure 1(a)(left). Moreover, there is no HRTO because consistent tradeups are found within its data (Figure 3): phage that are more infective to one bacterial host than other phage are also likely to infect more hosts. The bacteria-phage infection likelihoods can therefore be ordered, or graded. This grading can be quantified into a nested structure (Figure 2(b) gives statistics) which is significantly different from the modular structure that a lock and key would generate.

Now, the dynamics of graded, theoretical host-parasite systems differ from lock and key systems, as illustrated in Figure 1(b). Here, there is a Red Queen, as before in Figure 1(a), but the previously described game of hide and seek does not arise. Rather, hosts become more resistant through time and the parasites become more infectious, thus relatively little polymorphism is generated as hosts and parasites seek to optimise their respective functions. Bacteria seek receptor structures that no phage can bind, and as phage seek to bind even the most resistant bacteria, as a bi-product they can also bind susceptible bacteria better. Those susceptible bacteria, even if they appear as low-frequency mutants, are destroyed by phage and so this model, when accounting for a continual influx of new mutants, supports monomorphic clouds of phenotypes and little polymorphism (Figure 1(b,right)).

A molecular mechanism that could explain the nested Φ we observe would be where bacterial mutations controlled phage receptor expression levels whereby mutant bacteria expressed fewer receptors on their surface and so could gain resistance by loss of function mutations.^24, 25^ Phage could counteract this resistance mechanism by evolving greater adsorption affinities, allowing them to gain entry to all bacterial genotypes. Resistance under this scenario would confer a cost if the (potentially lost) receptor was important for growth, as it is in our system. Indeed, LamB is a transporter for maltodextrins which are supplied at growth limiting levels. In this case, receptor loss or down-regulation by mutation will increase phage resistance but it would reduce nutrient uptake and, therefore, growth rate, thus producing a CORT. While theoretically plausible, the latter does not arise in our model system. Instead, *E. coli* tended to evolve structural mutations in LamB (further discussion follows).

Nested interactions could also emerge from mutations if greater structural changes in the receptor, quantified by some geometric measure, conferred greater resistance - although such mutants would likely incur fitness costs by the impairing of LamB’s function as a nutrient transporter. However, while some of our library bacteria do pay these costs, we observed no CORT because certain bacteria have ‘Darwin monster’ phenotypes that bear no costs of resistance at all. Instead they derive growth benefits from their structural mutations (Figure 4(a)). Some library strains are even resistant to *all* library phage and yet grow more quickly than the ancestral strain in all six maltotriose concentrations used to assay growth; these Darwinian monster bacteria are named ‘26a’ and ‘56a’ (Figure 4).

**Figure 4:**
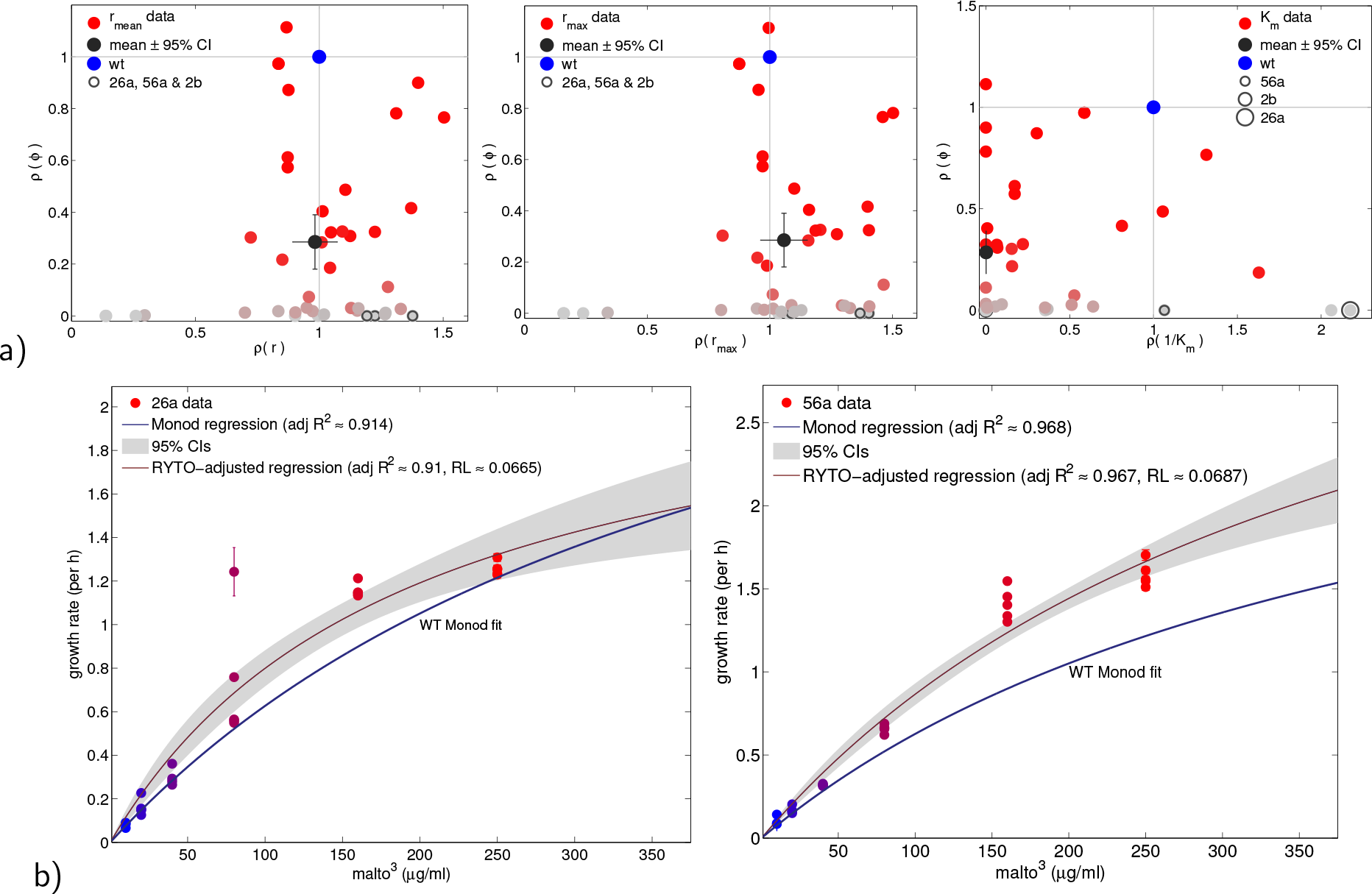
A reduction in phage susceptibility from wild-type increases growth rate in some strains, but decreases it in others. Using *ρ* here to represent the ratio of a phenotype with respect to the wild-type value, this shows mean phage susceptibility (*ϕ*) in the bacterial library is significantly lower than wild type. However, indicative of cost-free resistance, mean growth rates *r*_mean_ and *r*_max_ (as defined in Methods) show no reduction. Some strains exhibit higher than wild-type growth rates, including highlighted strains 2b, 26a and 56a that are pan-phage resistant. The mean half-saturation phenotype 1/*K*_*m*_ (averaged over 6 maltotriose supply concentrations) *is* significantly lower than the wild-type strain, indicating a significant increase in growth rate in some environments. b) Maltotriose concentration versus growth rate are shown for wild-type, 26a and 56a (nonlinear regression to data is shown as a line with grey confidence intervals, see Methods) indicating 26a and 56a exhibit faster growth than WT at *all* 6 maltotriose concentrations tested (error bars are 95% CI, *n* = 6 per condition); the latter are pan-phage-resistant strains. The WT nonlinear regression is shown here without error bars for clarity (*R*^2^ ≈ 0.997), for details on the WT data and model fit see Figure S7.

These observations lead to the question: what mechanisms of resistance to phage can simultaneously increase phage resistance and bacterial growth rates in a manner consistent with all of our data (Figures 2 - 4)?

Whole-genome sequencing of a subset of the bacterial strains indicates the bacteria evolved resistance in three membrane-related genes known to foster resistance mutations, *manYZ*, *malT* and *lamB*. Importantly, no genome possessed two resistance mutations, allowing us to interpret the effects of mutations on resistance (Table S1). Subsequent targeted Sanger sequencing of *lamB* for all bacterial isolates shows that the majority of strains have *lamB* mutations; 26a and 56a have nearby deletions in *lamB* (Table S2). Seeking structural LamB changes that could correlate with phage resistance and pan-resistance, we used a folding algorithm ‘Modeller’ (https://salilab.org/modeller/) to approximate mutant LamB structures. We then sought differences between the wild type LamB and the mutant to ascertain whether regions on LamB would correlate with the resistance phenotypes in Figure 2(a).

For this, we needed to identify differences in two protein structures of different sizes, so we developed a heuristic which is a fast extension of the Hungarian (aka *munkres*) matching algorithm (Supplementary 1.4) that compares sub-structures of the smaller protein to find optimal matches within the larger protein. We applied the algorithm, representing it as a mathematical function *M* (*p*; *q*) where *p* and *q* are protein structure vectors, to the wild type LamB alpha carbon (C*α*) structure, using it as a backbone against which other LamB structures were compared. The function *M* returns a vector in the lowest dimension of *p* and *q* that optimally matches a sub-structure of the higher-dimensional vector to the lower-dimensional vector with respect to Euclidean distance.

Now, the wild type ancestral LamB has 420, 3-dimensional C*α* coordinates (call them 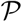) and the mutant named 26a has *m* = 421 coordinates (call them 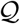), their algorithm-derived comparison 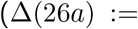 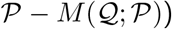 is illustrated in Figure 5. This image shows the 420 × 3-dimensional C*α*-difference vector Δ(26*a*) which is the difference of the matched protein subsets in (*x*, *y*, *z*)-space following convergence of the M-algorithm. This difference determines structural ‘hotspots’ which are defined to be those C*α* matches contributing more than *p*% of the total, between-protein difference in Euclidean space. Throughout, we made the arbitrary choice of *p* = 1 (Supplementary Text §1.4.1 gives details).

**Figure 5:**
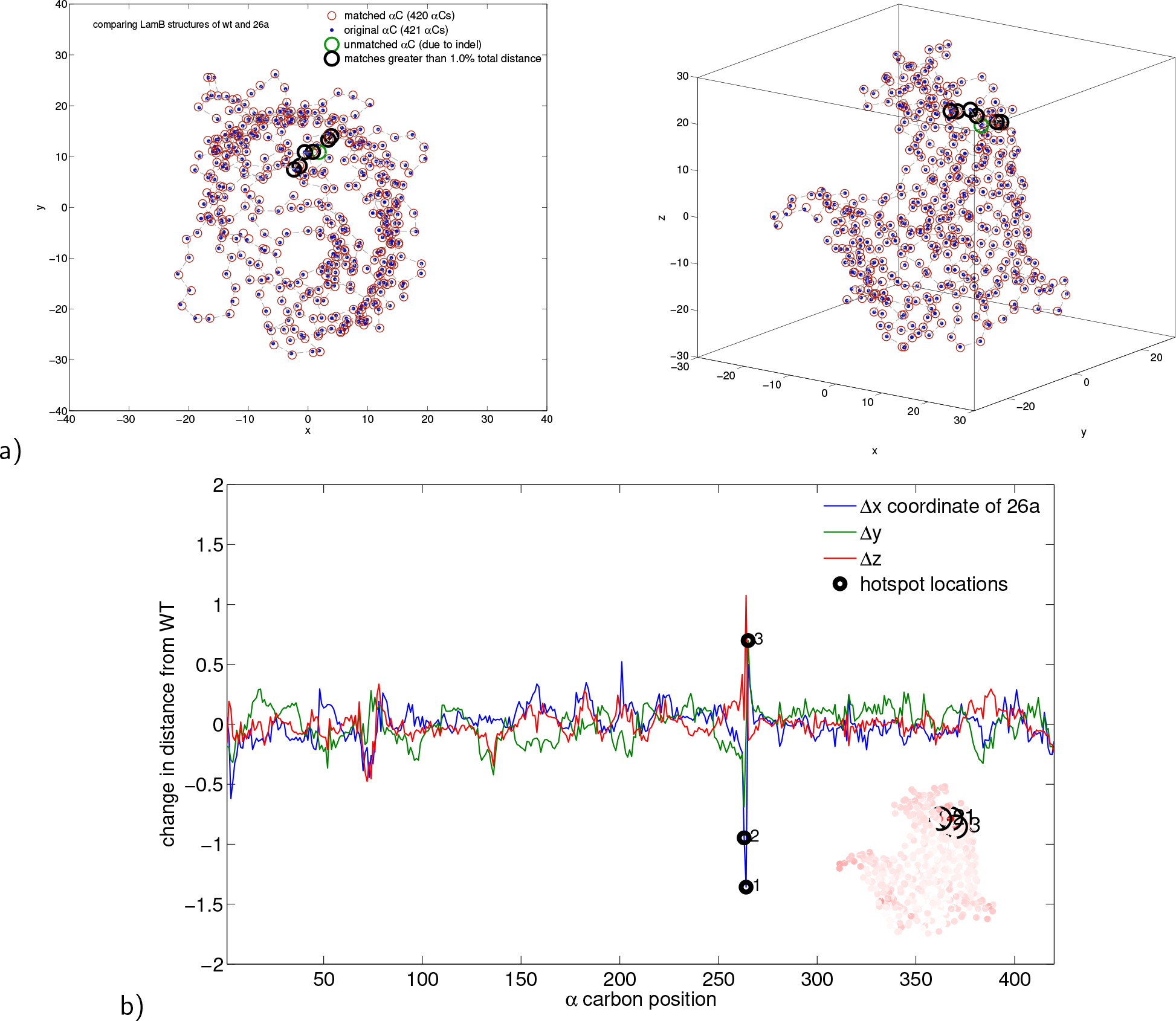
The bacterial genotype 26a that exhibits faster than wild type growth in all environments tested (Figure 4(b)) has a localised structural change in the phage receptor. A morphometric analysis shows that a deletion in *lamB* (see 26a in Table S2) results in a localised change in LamB that connects directly to the loops where phage *λ* binds. Plot a) superimposes 26a onto the WT LamB structure with the region of largest change (a ‘hotspot’) highlighted in black. Plot b) shows the 3d differences between WT and mutant LamB structures by unwrapping the entire protein loop into 3, one-dimensional timeseries wherein the hotspot can be seen as 3 spikes, one per physical dimension.

Using this definition, Figure 5 shows Δ(26*a*) has a localised structure for the pan-resistant strain 26a whereby three adjacent hotspots reside within the outer loops of LamB. Analogously, Figure S11 shows Δ(56*a*) has a hotspot in a similar location to 26a but with additional hotspots distributed across the protein. For comparison, Figure S10 shows Δ(2*b*) where 2b is another pan-resistant strain with a localised LamB change having two hotspots located at the entry point to the greasy slide.^26–29^ Figure 6 summarises the Δ(*m*) data for all mutants, *m*, that have over 400 alpha carbons which are shown as a red-greyscale heatmap superimposed upon the wild type LamB structure. Visually, this indicates redder regions (the hotspots) reside in both the outer loops and within the greasy slide.

**Figure 6:**
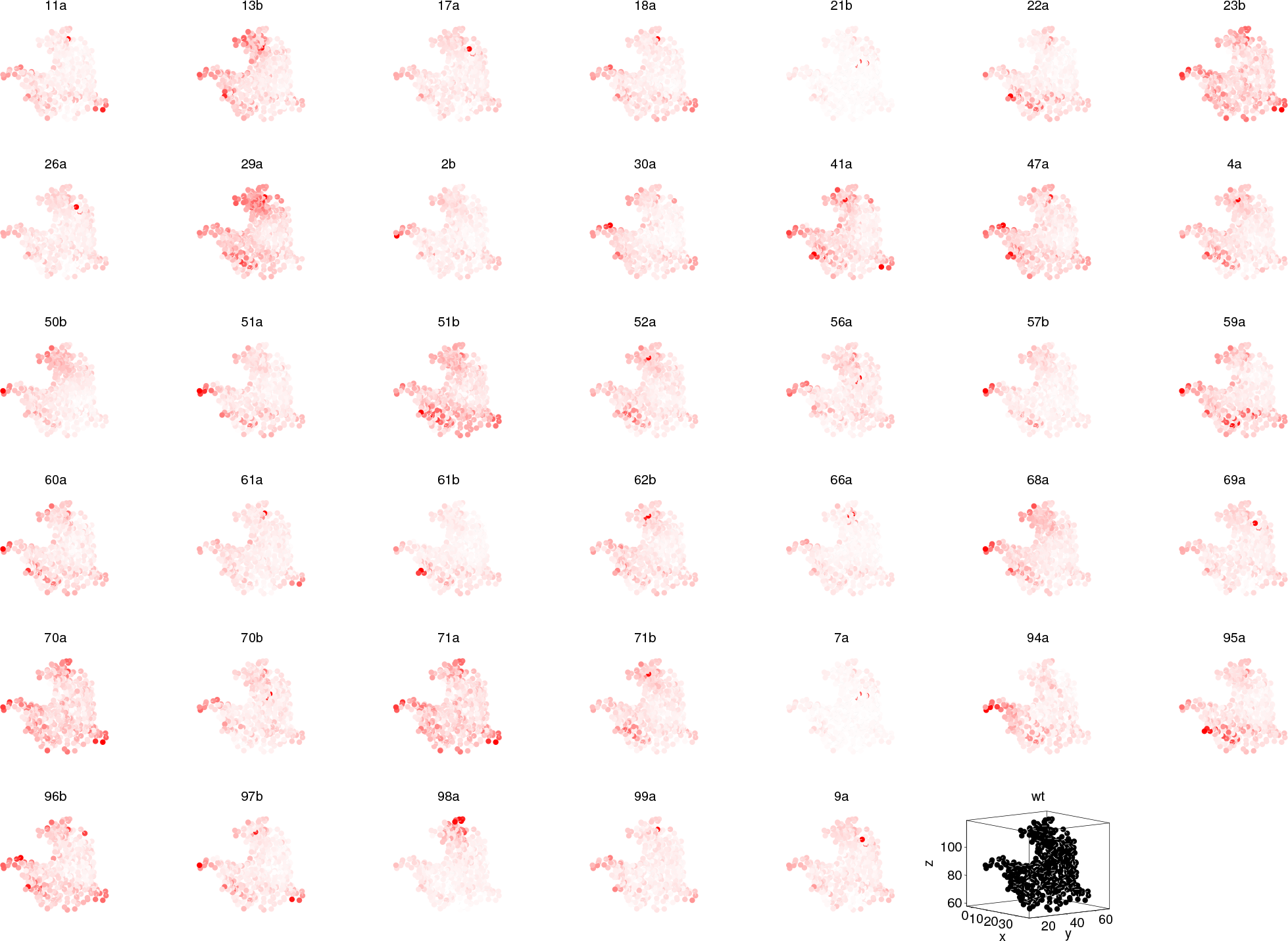
Structural LamB changes cluster in the greasy slide and the extracellular loops. This is a morphometric analysis of LamB structural changes from wild-type for all mutants in the bacterial library having 400, or more, alpha Carbons; these are shown as heat maps where white/grey represents no/little structural change and red is a high degree of change. This key is then superimposed upon the WT LamB structure for each mutant. The black LamB in the bottom-right corner is the wild-type 3d structure shown as a reference. Note how the reddest regions cluster around the greasy slide and the extracellular loops; Figure S12 uses a cluster analysis to demonstrate this quantitatively.

To quantify which LamB hotspots correlate most with adaption to phage and growth on maltotriose, we applied a *k*-means clustering analysis to the collated Δ(*m*) datasets, namely to ⋃_all *m*_ Δ(*m*). The number of clusters was determined from the local maxima of a kernel density estimate of the collated data (Figure S12(a)). The latter figure shows cluster centres located in LamB’s outer loops and other centres are located near the greasy slide. We undertook analogous clustering analyses for three bacterial sub-groupings, using (i) just library bacteria with LamB indels, (ii) just bacteria with LamB SNPs and (iii) just the pan-resistant mutants (a subset of the LamB indels grouping). The resulting clusters are shown in Figure S12(b). These 3 groupings all exhibit hotspot clusters in two biologically important regions: at the entry to the greasy slide and at the base of one outer loop; the indel group has hotspot clusters in the outer loops and while the SNPs group has hotspots in those loops, it has hotspots in the greasy slide too.

We then asked whether there was any correlation between distance in protein space from the wild type and increase in resistance to phage measured using the phenotypes from Figure 4(a). Figure 7(a) gives some, albeit weak, evidence for this by showing a significantly positive correlation between protein distance and change in infectivity using three different correlative measures. Thus the severity of LamB structural changes explain less than half of the variance in phage resistance and a circular dendrogram (Figure S6) corroborates this weak correlation visually. A rapid transition is apparent from the Hill function regression in Figure 7 whereby highly resistant bacteria have only slightly modified their LamB shape from wild-type.

**Figure 7:**
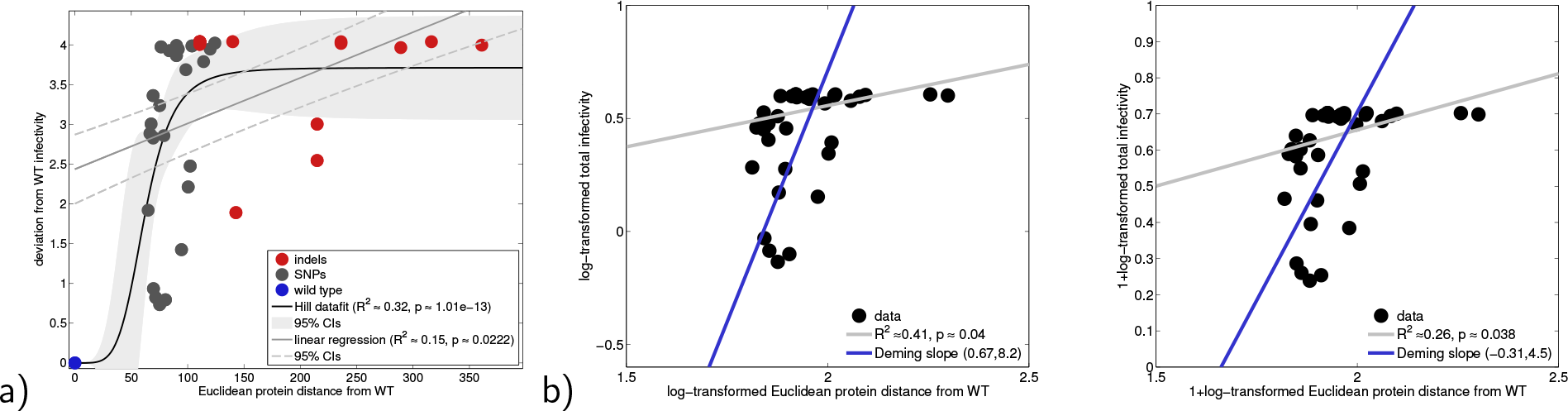
LamB protein distance-from-wild type correlates with change in phage susceptibility. Focussing on 41 LamB structures without large deletions (those with over 400 alpha carbons), a linear regression and a nonlinear Hill function regression show that Euclidean distance from wild-type in LamB shape space correlates significantly, if weakly, with Euclidean distance from wild-type’s phage susceptibility vector as determined using imaging algorithms (legends summarise regression statistics). b) Two linear regressions (standard in grey and Deming in blue) on log-transformed data from (a) show weak but significant correlations.

Given that LamB changes likely alter nutrient uptake, we asked whether a rate-yield tradeoff (RYTO) might be present in the strain library. Previously, we discovered RYTOs for the ancestral strain when it was grown at different resource concentrations:^3^ high resources yielded the fastest growth rates and less efficient growth. Here we tested whether the derived library strains also possessed within-strain and between-strain RYTOs. By relating growth rate and population density achieved per maltotriose supplied (*a.k.a.* biomass yield) for eight different maltotriose concentrations we consistently uncovered a within strain RYTO (Figure 8). However, between-strain data do not tradeoff: Figure 8(c) shows rate versus yield for each maltotriose supply concentration separately. These data are clear: in a fixed maltotriose environment, mutants strains with greater growth rates also have higher yields, thus we observe a between-strain rate-yield tradeup.^22^

**Figure 8:**
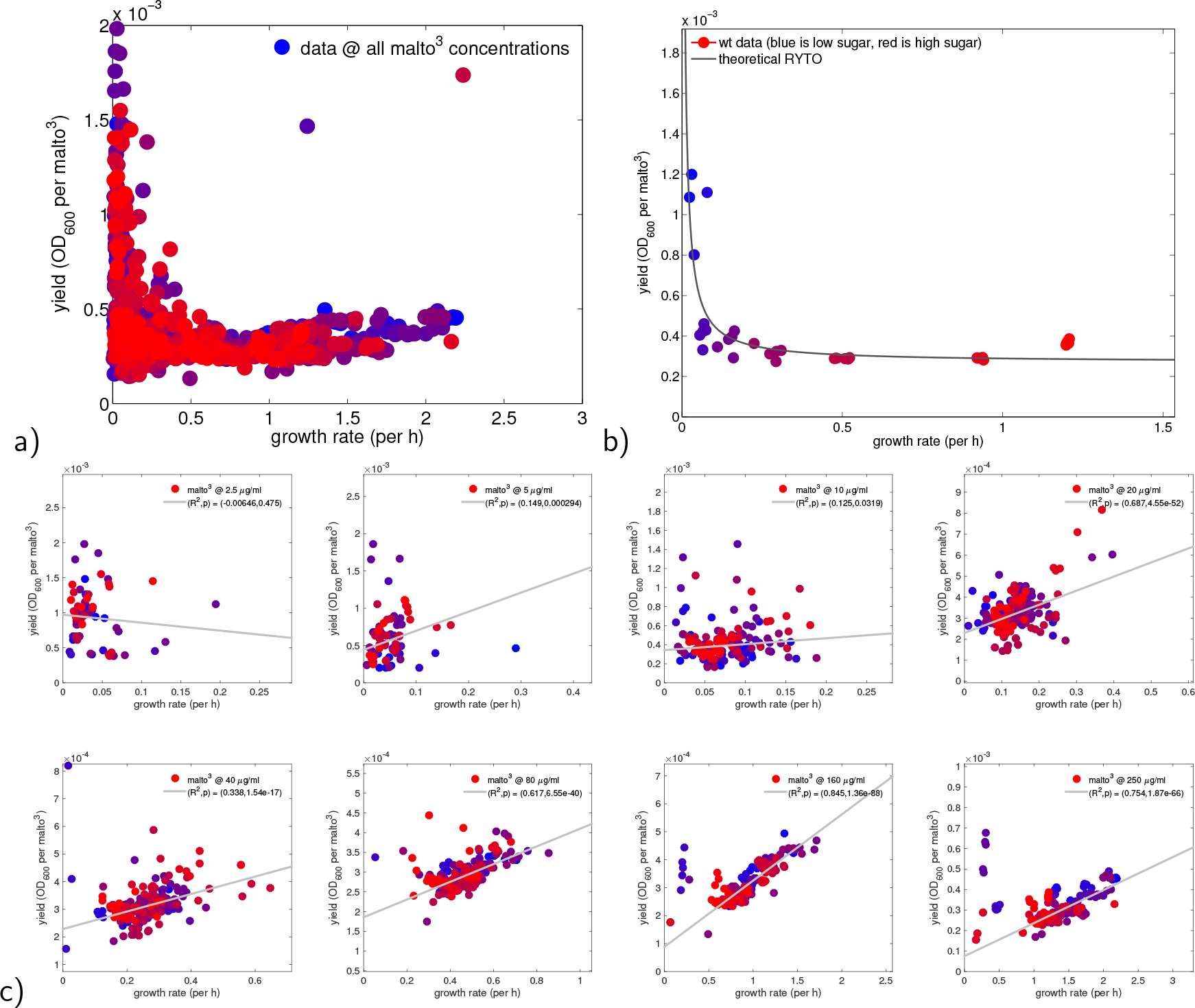
Bacteria exhibit rate-yield tradeoffs within-strain and rate-yield tradeups between-strain. a) This shows collated growth rate and yield data for all bacterial strains at all maltotriose concentrations tested; note how it appears to indicate a rate-yield tradeoff: at greater growth rates, biomass yield per maltotriose supplied is lower, creating an ‘L’ shape. But these data are hiding two conflicting trait relationships. To see this, notice how the yield data first decreases as rate increases, consistent with a tradeoff, but the yield data then increases with rate at higher values of the latter. This is indicative of a partial tradeup. (b) Investigating the latter in more detail, rate-yield data *within-strain*, here the wildtype, exhibit rate-yield tradeoffs for all but 2 strains (Figure S8 has data for all) which cannot explain the tradeup. (c) But *between-strain* rate-yield dataset can: significant positive correlations (tradeups) in 7/8 environments tested (p-values in legends), where correlations are greater at higher sugars.

There are at least two mechanisms by which mutations that increase phage resistance could also increase growth rate. The first is synergistic pleiotropy of that mutation: a structural change could simultaneously increase nutrient uptake rate and reduce binding affinity of the phage tail; this would increase growth rate. However, this cannot explain all our data: Figure 8(c) shows between-strain growth rate increases also increase yield. However, Figure S8 shows that growth rate increases correlate with decreased yield for most strains when more maltotriose is supplied to each genotype (this is the within-strain RYTO). Thus, the hypothetical synergistic pleiotropy that increases phage resistance, maltotriose uptake and therefore growth rate should, in the absence of other mutational changes, decrease yield and yet it does not. This synergistic pleiotropy is, therefore, unlikely to be the mechanism underpinning our data because the structural change in LamB must increase phage resistance, growth rate *and* yield.

The following argument, however, shows what can happen. The key idea is this: slower resource consumption due to a structural mutation that hampers phage binding could produce more efficient growth, and thus greater growth rates. To formalise this, let us represent growth rate, *G*(*S*), as a function of the extracellular sugar concentration (here maltotriose), *S*, by writing

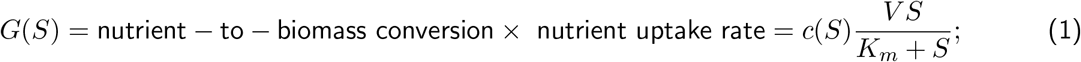

*V* is a mutable maximal sugar uptake rate phenotype, *K*_*m*_ is a half-saturation phenotype and *c*(*S*) is cell-yield measured per nutrient supplied, *S*. Nutrient uptake rate depends on both *S* and transport properties of the cell (*V* and *K*). From data, Figures 8(b) and S8 show that a decreasing function is a potential model for *c*(*S*) whereby

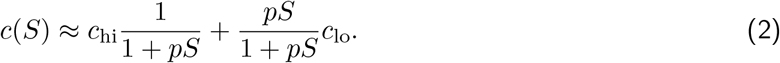

Biologically, *c* ought to be a function of the internal nutrient concentration, *S*_*i*_ say, and not *S*. But using an uptake equation of the form

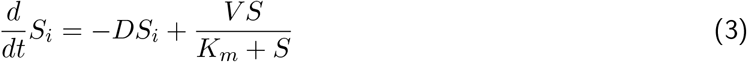

which we assume to be in equilibrium, we can relate *S*_*i*_ to *S*, namely 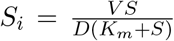, where *D* is the diffusion rate of the nutrient out of the cell. This relation can be re-arranged so that *S* = *S*(*V*, *S*_*i*_), ignoring the dependence on *K*_*m*_, at which point *c is* a function of the internal nutrient concentration. Note that *S* and *S*_*i*_ are positively correlated, as they should be.

Now suppose a mutation reduces the binding affinity of phage to LamB which, different to the previous argument, increases resistance but now instead leads to a decrease in the maximal uptake rate, *V*; we denote this small change in *V* by *dV*. This is reasonable because mutations that hamper phage binding could also impede nutrient uptake. Thus, uptake rates are reduced and our problem reduces to asking whether there are any circumstances under which *G*(*S*) increases when *V* decreases, or is 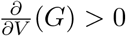 when *dV* < 0? To address this, elementary calculus tells us that

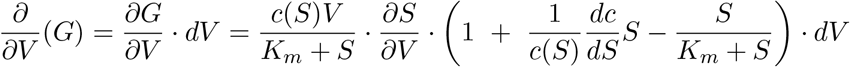

and this expression is positive, for negative *dV*, when the bracketed term is negative.

Now, the definition of a within-strain RYTO is that *dc/dS* < 0, and a short calculation shows that *G*(*S*) can increase following a transporter mutation with effect of size *dV* provided certain numerical conditions apply (Supplementary 5), one of which states that the maltotriose supply concentration must be sufficiently large. This is consistent with our rate-yield correlations (Figure 8(c)) which shows that rate only positively correlates significantly with yield between strains when maltotriose supply is greater than 2.5*μg/mL*.

We conclude that if the localised LamB phage-resistance mutations decrease sugar uptake rates and if the within-strain rate-yield tradeoff is strong enough, the growth rate reduction resulting from impaired uptake can be rescued by improving yield. Our argument here is that if phage resistance forces a reduction in nutrient uptake, then the increased yield from a RYTO can increase growth rate. Thus, just as two negatives multiply to form a positive, tradeoffs can be composed to give fitness advantages in the right circumstances.

Finally, we sought an additional tradeoffs by examining differences between lag times of susceptible and pan-resistant bacteria. We found weak evidence of this tradeoff whereby pan resistance incurred a small increase of about 1h in lag time for 2 out of 8 maltotriose concentrations tested (Figure S5).

## Discussion

Tradeoffs are usually studied in isolation and there are few datasets on multiple interacting tradeoffs. However, bacterial fitness is a complex function of many traits and since certain trait pairings tradeoff in isolation, because tradeoffs are not transitive elementary reasoning tells us those pairs must combine to form tradeups. Indeed, if trait X trades off with Y and trait Y trades off with Z then X and Z will tradeup (Supplementary §6). Thus, trait correlations can emerge in 3-trait datasets that are invisible to 2-trait datasets, thus it is unlikely tradeoff geometry can be safely inferred from such low-dimensional, 2-trait data.^4^

Given this, to better understand the processes by which bacteria resist phage, we examined several tradeoffs using data from protein to population scales on (i) whether phage mutants specialise, or generalise, on a library of bacterial mutants and (ii) whether bacterial fitness decreases as resistance to phage increases. Given the complexities, we delineate our discussion according to the four tradeoffs.

First, take the canonical host-range tradeoff (HRTO). This is the idea that parasites have limited choice, infect few hosts well or many hosts poorly. We found no evidence for this in phage *λ*, quite the contrary: our data exhibit tradeups whereby phage can be graded according to their infective ability (Figure 3).

Second, consider the cost of resistance tradeoff (CORT). The anticipated host costs for viral resistance in our data exhibit substantial variation. Some bacterial mutants pay a fitness cost for their phage resistance but others, quite unexpectedly, achieve fitness increases from phage resistance (Figure 4). In extreme cases, we even observe pan-resistant bacteria that have *improved* growth rates relative to the wild-type that are within approximately 15% of the fastest observed bacterial growth rates (Figure 4). In short, there is no CORT in our libraries.

Our argument explaining these ‘Darwinian monsters’ with multiple, simultaneous trait improvements is this. Structurally localised deletion mutations occur in the phage binding protein, LamB, that increase phage resistance by reducing binding affinity of the phage to LamB. It is unlikely, but possible, that these mutations also increase the uptake of maltodextrins into the cell, thus increasing growth rate and thus explaining cost-free resistance. However, our data are problematic for this idea as they also indicate the increased growth rates are due to increased yields. Now, increasing maltodextrin uptake does not explain this because a rate-yield tradeoff is present for most strains (Figure S8). Instead, it is likely that the localised phage-resistance mutations hamper the entry of maltodextrins into the cell, just as they hamper phage. However, the subsequent reduction in nutrient uptake can increase growth rates because the latter is composed of two processes in which tradeoffs are present: first the transportation of nutrients into the cell and, thereafter, the conversion of nutrients into biomass. Our theory shows that phage resistance and reduction in nutrient transportation by LamB mutation can, in fact, increase biomass conversion efficiency when a rate-yield tradeoff is present, and it is.

This motivates our third tradeoff, the rate-yield tradeoff (RYTO). Data show most cells harbour a RYTO whereby the metabolic efficiency of nutrient to biomass conversion decreases as nutrient supply increases. Thus, changes in growth rate arising from *lamB* mutations can be seen as a tensioning between two opposing forces: nutrient transport decreases, so intracellular sugars are less abundant (which could reduce growth rates), but metabolic efficiency could increase as a result (which would increase growth rates). The sizes of these respective increases and decreases will, in practise, dictate whether growth rate increases, or not, due to improvements in nutrient uptake and theory predicts a range of possibilities (Supplementary 5). This argument explains why there is no between-strain RYTO among the LamB mutants: strains that have increased growth rate have done so precisely because yield improved, thus yield and rate appear positively correlated in between-strain data (Figure 8).

We conclude that microbes operate in a complex, high-dimensional landscape of interacting phenotypic traits which might be at the biophysical, physiological and metabolic limits where tradeoffs are expected. Equally, they can operate in states free of such constraints and this might explain positively correlated adaptation in traits. But to explain our observations on growth improvements due to phage resistance mutations, our data needs the idea that tradeoffs can interact to produce correlated, pleiotropic benefits from a single mutation. This model has limited predictive power for understanding clinical infections and their evolutionary responses, but the idea that a cost-free resistance tradeup is a rational bacterial response to coevolution with phage within 3 days seems troubling from the perspective of mitigating resistance during phage therapy.

## Methods

### Bacteriology and phage assays

#### Phage and bacterial mutant library generation

We co-cultured *E. coli* strain B REL606 with the obligately lytic *λ* strain cI26. When co-cultured with cI26, REL606 experiences pressure to evolve resistance because *E. coli B* strains lack generalized phage defenses such as mucoid cell formation, restriction modification, or cRISPR adaptive immunity.^30, 31^ This pressure is magnified by the lytic phage’s increased virulence as compared to its lysogenic relatives.^32^ Once *E. coli* resistance evolved, phage experience selection to evolve counter-defenses, triggering a rapid arms race within days.

100 flasks were initiated with few phage (~ 10^2^ particles), and 10^3^ bacterial cells that were preconditioned in the experimental environment for 24 hours. Small initial populations increased the likelihood that mutations for defense and counter-defense arose *de novo*, which improved our chances of isolating unique *lamB* mutations and evolving divergent phage genotypes between replicates. The flasks were filled with 10ml of modified Davis Medium (DM)^33^ (125 *μg/ml* maltotriose instead of glucose and 1*μg/ml* of magnesium sulphate). Bacteria and phage were allowed to reproduce for 24 hours, at 37°C, and shaken at 120 rpm. At 24 hours, a random 100 *μl* sample of each flask was added to a fresh flask and the bacteria and phage were allowed to grow again. This cycle was repeated once more and phage and bacteria were sampled after the third day of growth. We ended the experiment at this early time-point to ensure that the bacteria could only acquire a single mutation for defence, which made linking genotype to phenotype more tractable. Additionally, in previous experiments we observed the greatest genotypic diversity of bacteria and phage on the third day (unpublished data). *E. coli* evolve resistance in this environment through many loci, however the most common is *lamB* (unpublished data). For this study we will focus just on the *lamB* mutations.

#### Bacterial and phage isolation and storage procedures

To isolate bacteria we streaked a sample on Luria Bertani (LB) agar plates,^34^ randomly picked two colonies and then re-plated two more times to remove all phage. Finally, we grew each colony in liquid Luria Bertani media overnight and preserved two 1 ml samples in 15 percent glycerol and frozen at −80°C. The entire phage population was preserved for each flask by chloroform preparation of the remaining volume of culture, 8ml.^35^ Clonal isolates of the phage were created by picking plaques (miniature epidemics derived from a single phage particle) from bacterial lawns (films of bacteria immobilised in soft agar spread on top of Petri dishes) (Adams 1959). For each phage population, we attempted to isolate phage from three separate lawns, one derived from the ancestral bacteria (REL606) and the two bacteria isolated from the very same flask. By using the coevolved bacteria for phage isolation we increased the chance of sampling phage that had evolved specialised interactions with their coevolved bacteria, thereby improving our chances at uncovering more phage diversity with modular bacteria-phage interactions than if we had only used the ancestral bacteria. Two plaques from each flask were isolated and clonal cultures were created according to Adams, 1959.^35^ When choosing phage, plaques were favoured that formed on the lawns of the coevolved bacteria.

#### Estimating E. coli resistance to a library of phage

We measured resistance by challenging each bacterium with every phage isolate. To do this, we made spot plates by dripping ~ 2.5*μl* of each phage stock on bacterial lawns.^35^ After 24 hours of growth at 37 degrees C the bacterial lawn would thicken, unless a phage was able to kill it, in this case a round clearing (spot) formed under the drip. Digital pictures were taken of each plate using an AlphaImagerTM 2200 by Alpha Innotech. The matrix of interactions was previously reported, however, clearing were determined by eye and only a ‘0’ or ‘1’ was recorded for whether or not a given phage infected a given bacterium.^36^

#### Competitive ability assays

Competitive fitness for lamB mutants were measured by competing each genotype head-to-head with a genetically marked version of the ancestor, REL607. REL607 can metabolize arabinose, whereas REL606 cannot because of a single nucleotide substitution that has little effect on the bacterium?s fitness. Marked and unmarked genotypes can be distinguished on tetrazolium arabinose plates, which provides a tool to estimate the relative frequency of each in a mixed population. Full descriptions for competition experiments can be found.^37, 38^ In short, we initiated each flask with 50% of the resistant type and 50% of its ara+ ancestor, cultured them for three days identically to the experiment except without phage, measured their initial and final densities and calculated the ratio of the Malthusian parameters for the evolved versus the ancestor. We performed three replicates for each lamB genotype.

#### Sequencing

. We sequenced *lamB* for at least one bacterial isolate from each flask, and the second isolate was sequenced if preliminary tests revealed the two sympatric isolates had different levels of resistance. Sequencing was performed with an automated ABI sequencer maintained at Michigan State University Research Technology Support Facility. PCR amplified fragments purified with GFX columns were used as templates. Fragments were amplified with primer sequences 5’-TTCCCGGTAATGTGGAGATGC-3’ and 5’-AATGTTTGCCGGGACGCTGTA-3’, placed 1,398 bases upstream and 504 bases downstream of the gene, respectively.

Full genomes of 15 *E. coli lamB* mutants were sequenced to determine whether any other mutations occurred in the *E. coli* genomes that may influence resistance. Isolates were chosen that we suspected had multiple *λ*-resistant mutations because they evolved high levels of resistance despite only possessing a single amino acid change in LamB, because a mutation in *lamB* was not discovered, or because the genotype possessed a distinct resistance profile. Technicians at the Research Technology Support Facility at Michigan State University sequenced the genomes using an Illumina Genome Analyzer IIx. Genomic DNA samples were created by reviving frozen bacteria in LB medium, growing them overnight and then isolating DNA from several millilitres of the culture with Quiagen genome tips. Samples were fragmented by sonication, prepared with barcoded attachments and run as multiplexed samples over four lanes. Mutations were predicted from the resulting 75-base DNA single end reads using breseq v0.13 (https://www.sanger.ac.uk/science/tools/ssaha2-0) and the ancestral genome (Genbank accession: NC 012967.1) used as the reference.

#### Protein shape prediction from amino acid sequence

Mutant protein shapes were predicted using the program Modeller (http://salilab.org/modeller/). Predictions were made in two steps: First, homology-based techniques were used to generate a protein structure by using the known ancestral LamB shape as a guide. LamB structure was determined by x-ray crystallography to 3.10-angstrom resolution (Genbank accession: YP 003047080) (Schirmer et al. 1995). Next, *de novo* loop refinement reorganized protein conformation to minimize entropy created by electrostatic conflicts introduced by the substituted amino acid.

### Theoretical computations and statistical concepts

All computer codes were implemented in Matlab, Java and ImageJ and can be obtained from the authors on request. Some can be downloaded from websites as detailed in the supplementary information.

#### Protein structure differentiation

To differentiate protein structures geometrically we implemented an iterative algorithm in Matlab (provided as executable code in Supplementary §1.4) that can be applied to two arbitrary, 3-dimensional datasets. We applied it to the alpha carbon (C*α*) coordinates of each protein determined by the above folding algorithm. Our bespoke matching algorithm seeks to match one C*α* dataset against another by performing rigid transformations that minimise the Euclidean distance between the two. Where C*α* datasets differ in size, due to an indel, the algorithm attempts to place the smaller dataset into the larger one iteratively, by deciding which points to ‘keep’ and which to ‘reject’ in the matching process so that the Euclidean distance between a sub-structure of the larger dataset can be placed as close as possible to the smaller one. There can be non-unique ways of making this choice (for example, delete the vertex of one cube and seek to match the resulting structure back into the cube) and a limitation of the algorithm is it only returns one possible coordinate match.

Figure 5 is one such matching where one structure is the C*α* backbone of the wild-type LamB maltoporin and the other structure is a deletion mutant named ‘26a’ which, as a result of the deletion, has a shorter LamB coordinate set. The C*α*s that cannot be matched as a result are shown as green circles in the figure, whereas one structure is shown with blue dots and the other with red rings encircling those dots.

#### Imaging bacteria-phage infectivity parameters

As detailed in Supplementary §1.3, imaging algorithms were implemented in Java / ImageJ to determine the size of a phage clearing (a ‘plaque’) on a bacterial lawn using images captured from experimental spot assays, Figure S1 shows one of these. This ‘size’ parameter was normalised against the largest observed size of all such assays for all bacteria-phage pairings and this was used as the value of the infectivity parameter for that phage-bacteria combination. The values resulting from this set of assays can be seen in Figure 2, an important region in which is the large black zone at the top of the matrix in this figure. This shows a cluster of bacteria that can be infected by none of the phage, these are the pan-phage-resistant genotypes.

### Coevolutionary dynamics: a mathematical bacteria-phage coevolution model

A system of partial different equations were used to produce the short-term coevolutionary dynamics in Figure 1 which are these:

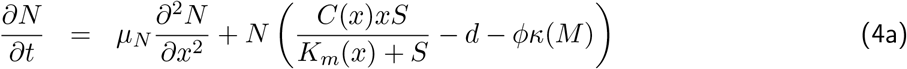

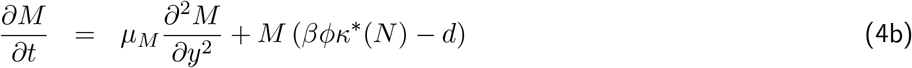

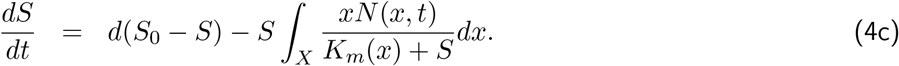

Here, modelled phage and bacteria coevolve in a chemostat where *N* (*x, t*) is the density of bacteria per unit volume at time *t* and *M* is the density of phage; the chemostat turnover parameter is *d* and the limiting sugar supply parameter is *S*_0_. The variable *x* ∈ [0, 1] =: *X* is a phenotype that controls several properties of the bacteria: first, the maximal uptake rate of extracellular sugars bacteria is equal to *x*, *C*(*x*) is bacterial cell yield measured in cells produced per unit sugar, *S*, then *K*_*m*_(*x*) is the half-saturation constant associated with cells of phenotype *x*. This formulation allows the model to encompass a range of tradeoffs, like the rate-yield and rate affinity tradeoffs, if need be.

Changes in *x* are modeled according to a linear diffusion process, with diffusion coefficient *μ*_*N*_, which represents the change in bacterial phenotype brought about through evolutionary mutations. There are many other ways of representing evolution in this framework, we use the one presented here for simplicity of presentation, not necessarily biological realism. For instance, this model allows the phenotype to evolve even when *C* = 0 and so no cell divisions are taking place. (This can be fixed, but at the cost of complicating the model.)

The diffusing evolutionary phenotype *y* ∈ [0, 1] =: *Y* associated with phage, of density *M* (*t, y*) at time *t*, is a recognition phenotype which is used to decide which of the bacterial phenotypes each phage, with burst size *β*, can successfully infect. The infection process is modeled through the pair of linear operators *K* and its adjoint operator *K*\*. The general structure for this operator pairing follows the reasoning that a phage of type *y* can successfully infect a bacterium of type *x* on contact with probability *p*(*x*, *y*). Therefore, assuming a mass action law of contacts and counting bacterial losses, we use a bacterial loss / death term of the form −*ϕ* ∫_*X*_ *p*(*x*, *y*)*N*(*t, x*)*M*(*t, y*)*dx*, where *ϕ* is a parameter that represents how frequent those bacteria-phage contacts are; for notational convenience we therefore define

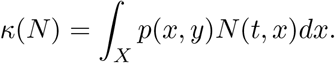

When we count (integrate over) that same number of bacterial losses from a phage perspective, which are not losses now but gains, we arrive at the integral

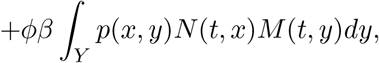

or

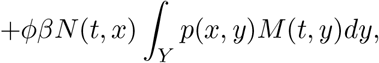

which involves an expression for the adjoint operator of *K*, namely

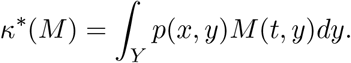

The nature of the infection is summarised by the detailed form of the function *p*(*x*, *y*). For instance, there could be said to be a lock and key interaction if, for each fixed value of *x*, there is a unique *y* that maximises the function *y* ↦ *p*(*x*, *y*) and vice-verse. A graded interaction would be one whereby for each *x*, infection likelihood increases with *y*, meaning *∂p*/*∂y* > 0 everywhere. The two ‘lock and key’ and ‘graded’ conditions described here are mutually exclusive.

For the simulations presented herein, we arbitrarily chose *p*(*x*, *y*) = *ϕ*(*x* − *y*) where *ϕ* is a Gaussian function in order to model a lock and key structure, we then used *p*(*x*, *y*) = *ψ*(*x* − *y*) where *ψ* is an error function in order to model a graded structure. The two heatmaps in Figure 1 illustrate these two choices of mathematical infection mechanism.

Finally, this mathematical model has a limitation common in the tradeoff literature as it represents two one-dimensional tradeoffs through relationships between the trait *x* (normalised maximal resource uptake rate) and the functions *K*_*m*_(*x*) (half-saturation) and *C*(*x*) (biomass yield). Thus the entire tradeoff geometry here is a curve (*x*, *K*_*m*_(*x*), *C*(*x*)) in a 3-dimensional space. In keeping with the remainder of this article, we could have modelled bacteria-phage coevolution by allowing (*x*, *K*_*m*_, *C*) to be a single, 3-dimensional evolving trait which would provide much greater ‘freedom’ to the evolutionary trajectories of these parameters, but at the expense of greater computational difficulties.

### Bacterial growth phenotypes

To quantify relationships in datasets between growth rate and phage resistance, we applied a Michaelis-Menten-Monod datafit to filter noise from all empirical growth rate data. Thus, for each bacterial genotype and each maltotriose supply concentration, we produced a mathematical curve representing growth rate parameterised by two phenotypes, *K*_*m*_ and *r*_max_:

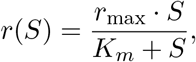

where *S* is the concentration of maltotriose used. Figures 4(b) show these datafits for two strains, comparing it to the wild-type datafit.

If we define *r*_mean_ to be the average growth rate observed over a range of maltotriose concentrations ranging from zero to *S*\**μg/ml*,

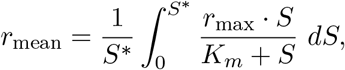

then *r*(*S*) and *r*_mean_ will increase if *r*_max_ increases or *K*_*m*_ decreases. These measures are used in Figure 4(a) to compare growth rates between strains where *S*\* is 250*μg/ml*.

It is standard to factor the dependence of growth rate in terms of uptake and metabolic parameters by writing *r*_*max*_ = *V*_*max*_ × *c* where *V*_*max*_ is the maximal nutrient transportation rate and *c* is cell yield per nutrient supplied,^39^ from where

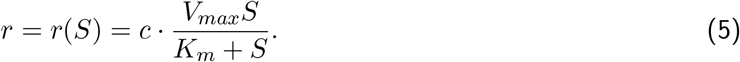

Under the circumstances where a rate-yield tradeoff applies, *c* need not be constant and it has been shown,^40^ to approximation, that the following model can be a better descriptor than a constant:

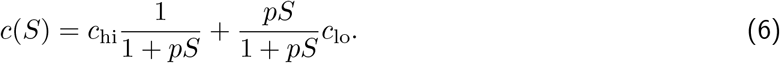

Putting these expressions together, a model for growth rate consistent with a RYTO is given by the following expression

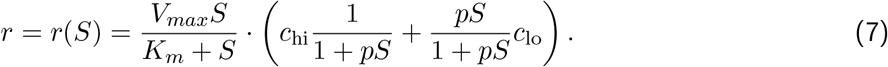

In the main text, we then argue that *c*(*S*) should depend on intracellular nutrients, not extracellular ones, so we introduce an equilibrium relationship between the two to account for this, equation (3).

As *r*_*max*_ = *V_max_ × c*, these parameters are not independent and they must be inferred from different data sets. We do this, for example in Figure 8, following previous analyses^22^ by first estimating population size as the carrying capacity, *K* (which, we note, is *not K*_*m*_), in the following logistic equation that is also parameterised by exponential growth rate (*r*):

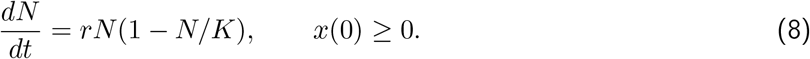

Solutions of this equation are modified to account for a lag phase (*L*) so that parameters *r*, *K* and *L* are determined from the best fit of the following 4-parameter model to microbial population density timeseries:

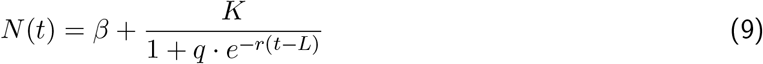

where *N* (*t*) is density at time *t*, *β* is a non-biological ‘blank’ parameter (it denotes the effective value of zero population density in a translucent microtitre plate) and *q* is a composite parameter that we do not make use of in our analysis. The value of *c* is then defined to be *K* divided by the maltotriose concentration supplied to the experimental device, which is known by design.

This procedure was repeated using bacterial density timeseries at 8 maltotriose concentrations ranging up to 250*μg/ml* which produces a dataset of yield values that are fitted using a model of the form (6). This procedure provides data on the dependence of *r* on *S* to which equation (7) is then fitted. This procedure produces the rate, yield and *K*_*m*_ data shown, for example, in Figures 4 and 8.

## Article and author information

**JM Contribution**: resources, funding acquisition, conceptualisation, data curation, formal analysis, validation, investigation, visualisation, methodology, writing – original draft, project administration, writing – review and editing ; **Competing interests**: no competing interests declared

**REB Contribution**: resources, supervision, funding acquisition, conceptualisation, data curation, formal analysis, validation, investigation, visualisation, methodology, writing – original draft, project administration, writing – review and editing, software ; **Competing interests**: no competing interests declared

**IG Contribution**: resources, supervision, funding acquisition, conceptualisation, validation, investigation, project administration, writing – review and editing ; **Competing interests**: no competing interests declared

**RPM Contribution**: conceptualisation, validation, investigation, software, formal analysis, methodology ; **Competing interests**: no competing interests declared

**MH Contribution**: conceptualisation, validation, investigation, methodology ; **Competing interests**: no competing interests declared

**CR Contribution**: conceptualisation, validation, investigation, methodology ; **Competing interests**: no competing interests declared

## 1 Supplementary Methods

### 1.1 Coevolution Mathematical Model Simulations

The Matlab codes used to simulate equations (4a-c) from the main text can be downloaded from https://github.com/rebear217/PhageMFiles

Upon downloading, the code produces Figure 1 on typing the command Go into the Matlab command window.

In case of problems, these files are mirrored at http://people.exeter.ac.uk/reb217/rebHomePage/data.html

### 1.2 Additional lag phenotypes

A measure of lag time *L*, as described in methods and derived from a mathematical model is used in this article. For completeness and to validate the robustness of any statements made regarding the lag phenotype, an additional measure of lag is used that is determined solely from a population density time series, thus avoiding mathematical model datafits.

The measure is this: given a series of times, (*t*_*j*_) and population densities, *N*_*j*_, lag time is estimated from the smallest index *J* for which a linear regression, *N* = *α* + *t* · *β*, of (*t*_1_, *t*_2_, …, *t*_*J*_) and (*N*_1_, *N*_2_, …, *N*_*J*_) returns a significantly non-zero value for *β* using an F-test. This test requires an acceptable slope parameter and a p-value associated with the regression which tests whether, or not, a slope of that size has been breached by data. If so, the algorithm accepts the claim the data is no longer constant at that time and the population is, therefore, increasing in size.

A stronger test was also implemented whereby the largest *J* for which all regressions with *j* ≥ *J* return significantly non-zero values for *β*, whereupon lag time is *t*_*J*_. The latter is less sensitive to fluctuations in data that could indicate apparent population growth, only for it to be the result of a temporary density measurement fluctuation. The claims we make in the article and supplementary regarding lag do not depend on which of these three possible lag statistics are used.

As the above algorithms are not commonly used in the literature, we include the following Matlab code which returns the index J (calling it `lagPointData.J`) where the variable `Nj` represents the times eries (*N*_*j*_). Note, the latter needs to contain at least 3 datapoints to determine J.

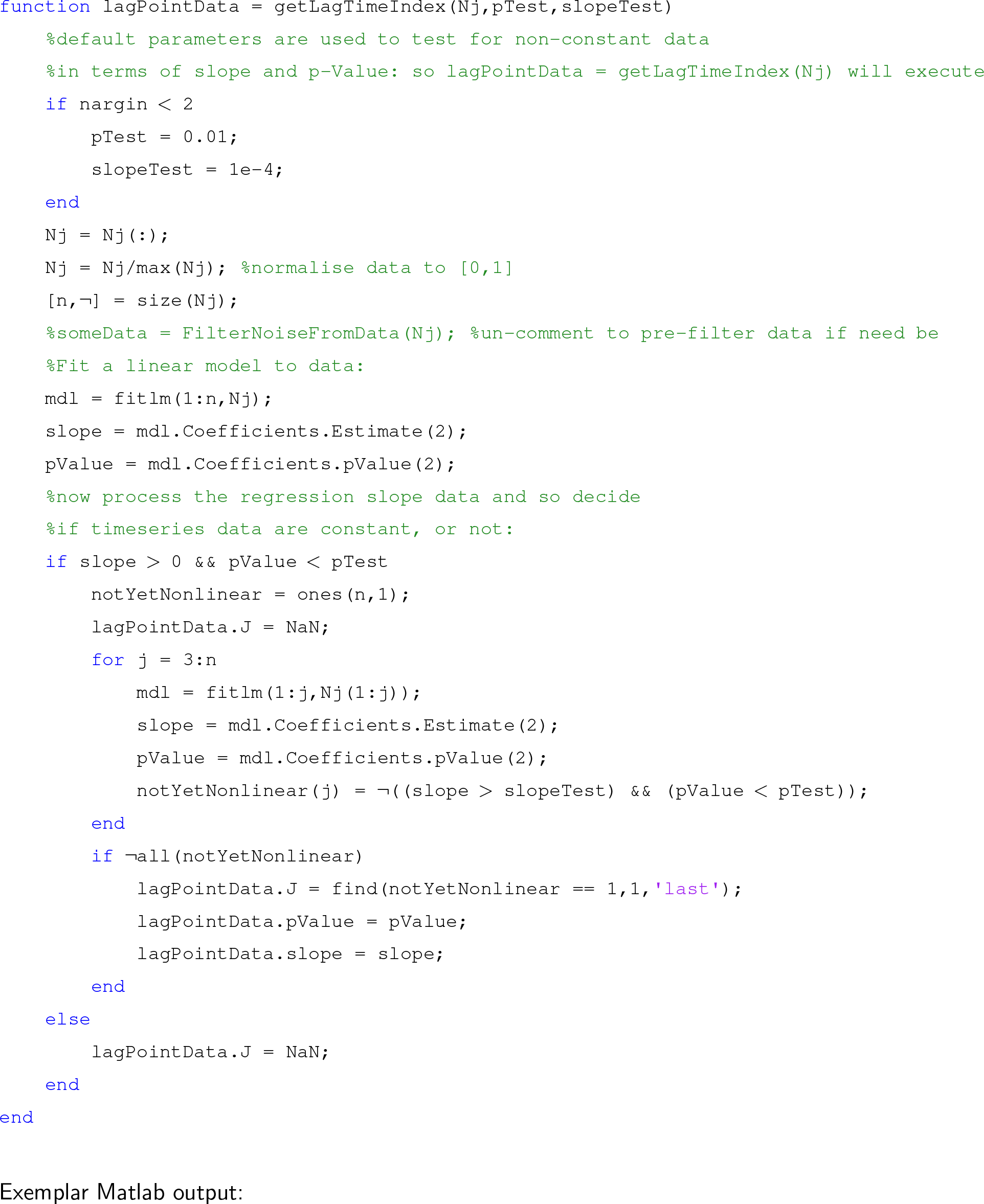

Exemplar Matlab output:

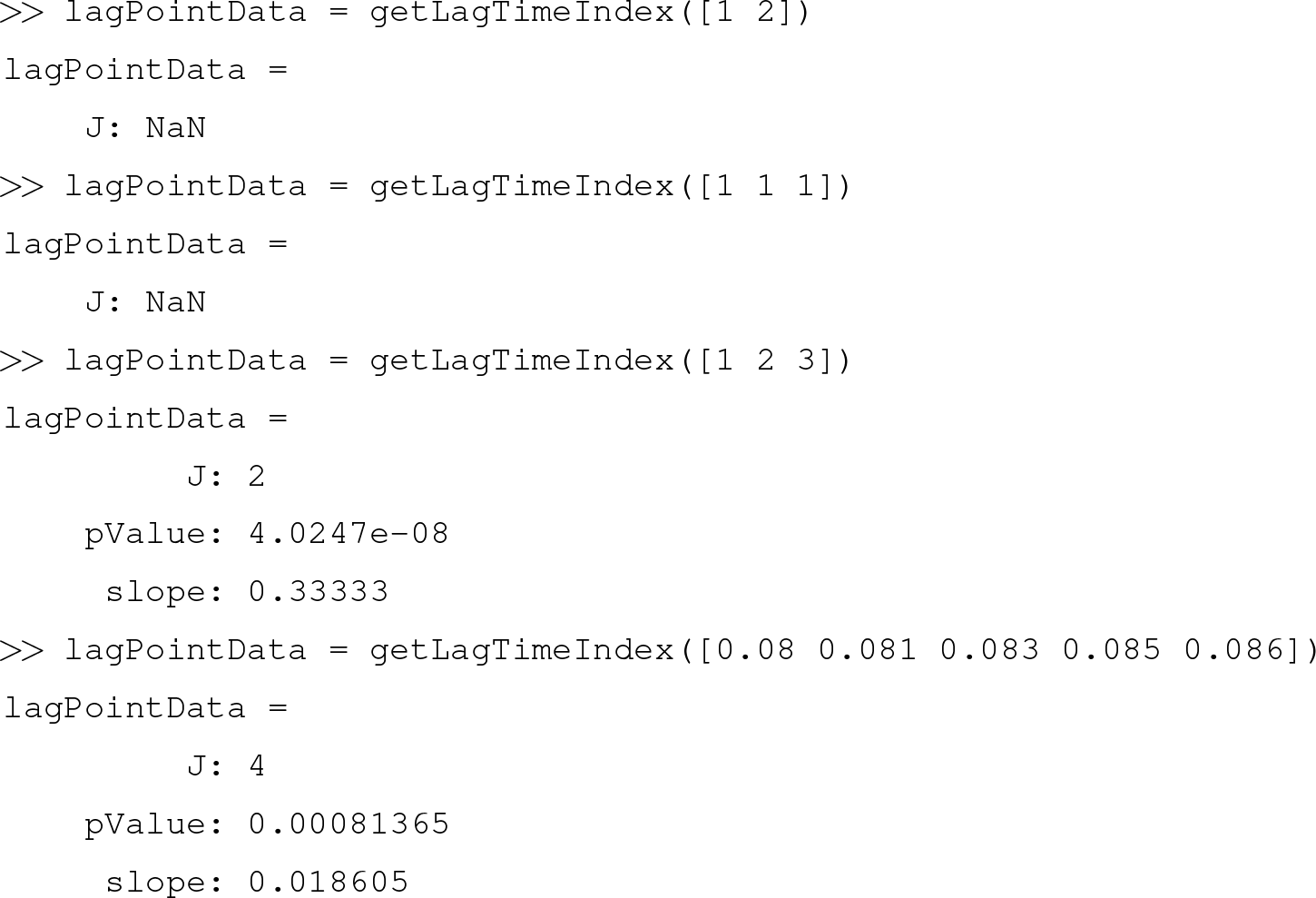

### 1.3 The Infectivity Matrix

Imaging algorithms used to determine the infectivity matrix in Figure 3(a) were implemented in Java and ImageJ. The methodology that determined this matrix is depicted in Figure S1 and the necessary codes and plugins needed for performing this analysis can be obtained from the authors.

Figure S1(a) shows a bacterial lawn with phage plaques. To determine infectivities for phage and bacterial susceptibilities, bacterial lawn images were analysed quantitatively to determine the clearing size based on a thresholding criterion, an example of which can be seen in Figures S1(b) and (c). Figure S1(d) then shows a colour-coded infectivity vector that results from this process for each phage with respect to the bacterial genotype cultured on this particular lawn. Phage infectivities and bacterial susceptibilities are given in arbitrary units and take values ranging upwards from phage resistance (a value of zero where there is no plaque) to higher values which are limited by the image colour representation of the photograph image.

### 1.4 LamB Morphometrics

We now describe the algorithm we implemented that attempts to make a comparison of two sets of *N*-dimensional vectors in two different Euclidean spaces of arbitrary dimension, here *N* = 3 for the protein comparison applications described in the main text. Matlab implementations of the algorithms can be downloaded from https://github.com/rebear217/PhageMFiles which, as an example of how the codes execute, produces Figure S2 on typing the command sphereTest into the Matlab command window.

**Figure S1:**
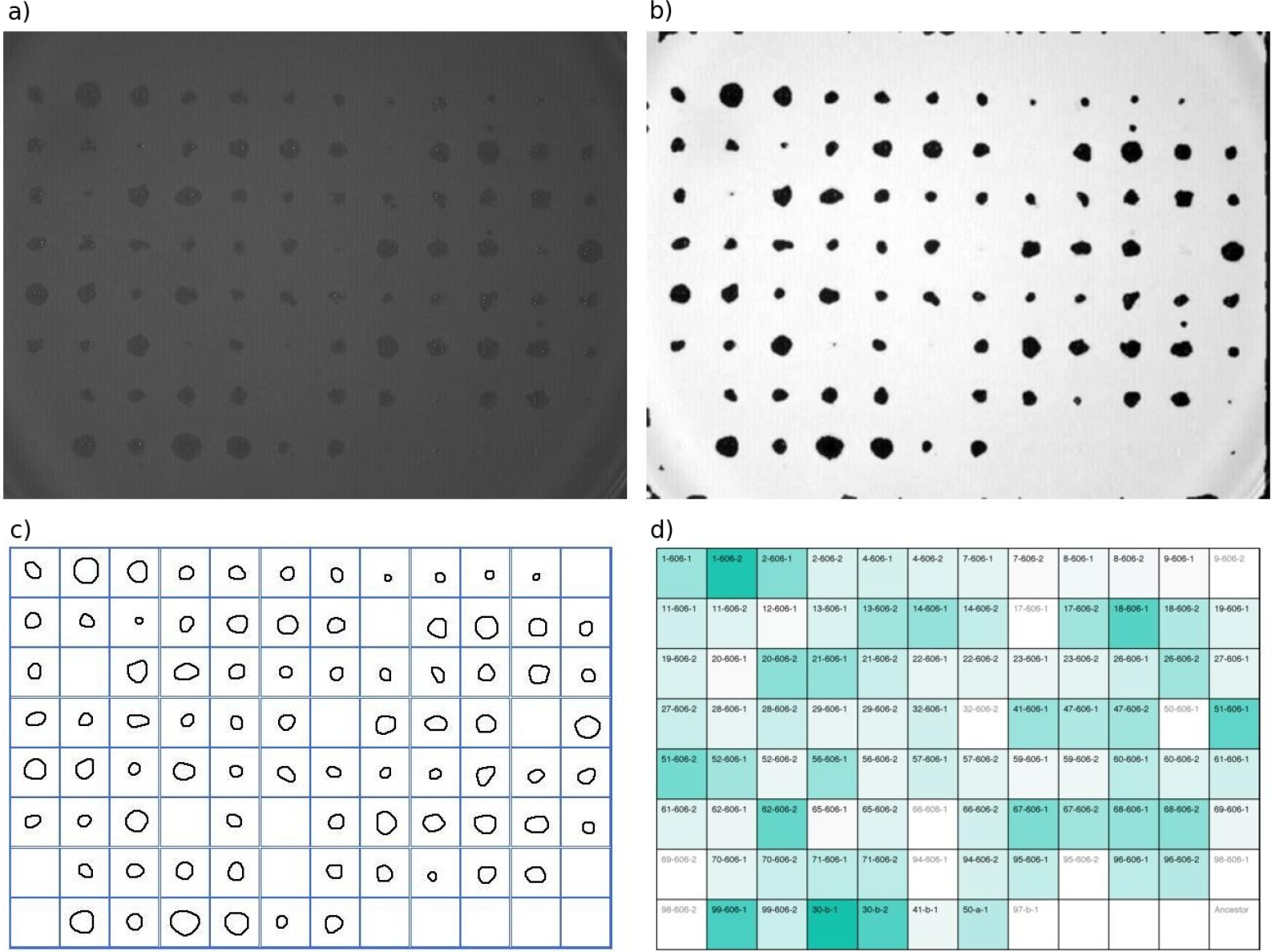
An exemplar assay for a lawn of bacteria named ‘23B’ tested against all library phage and so illustrating the image processing steps needed to quantify phage infectivity patterns from the image of a plate. a) A photograph of the lawn taken at 72dpi showing plaques produced by all the phage. b) A pre-processed image shows the plaque blob geometry to be quantified. c) A blob-detection algorithm further simplifies the plaque data. d) The resulting normalized infectivity data is shown as a green heat map matrix where darker green denotes higher likelihood of infection. The data from (d), which is a series of values between zero and one, are then collated into a row vector that will be subsequently placed into a matrix form for further analysis (for example, placed into a matrix like the one shown in Figure 2).

Note that our codes makes use of the following 3rd-party Matlab code ‘Functions for the rectangular assignment problem’ by Markus Buehren: https://uk.mathworks.com/matlabcentral/fileexchange/6543-functions-for-the-rectangular-assignment-problem which needs to be installed in order for our exemplar test codes and algorithms to work.

**Figure S2:**
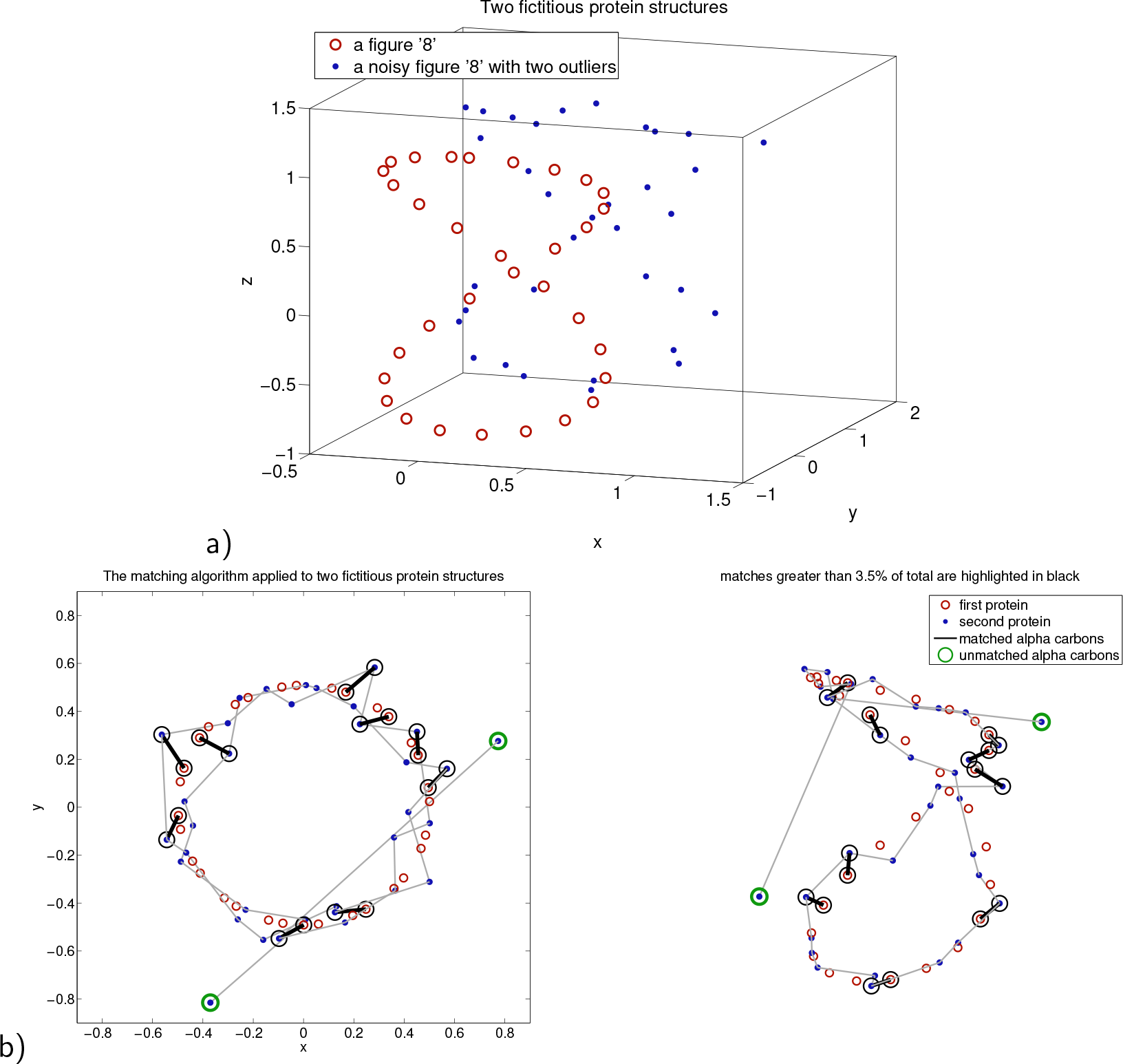
a) We present some synthetic data to illustrate the iterative protein matching algorithm, as follows: first, a figure ‘8’ is embedded in 3d space (32 red circles) which is then perturbed with noise and two points are added to it (34 blue dots). If the former dataset is thought of as representing the *α* carbon locations of a certain protein, the latter can be thought of as a mutant structure with an insertion mutation of two *α* carbons. b) The result of the algorithm is an optimal match between the original figure 8 and a 32-point subset of the noisy figure 8. The remaining, unmatched two points determined by the algorithm are shown in green. Certain matching pairs on both figure 8s are indicate by thick black lines where 3% of the total Euclidean distance between the matched test figures is due to that one particular pairing; we call these pairs ‘3% hotspots’; *p*% hotspots are analogously defined, smaller *p* values have fewer hotspots. The grey lines guide the eye through all the blue-dotted ‘protein’ points.

**Figure S3:**
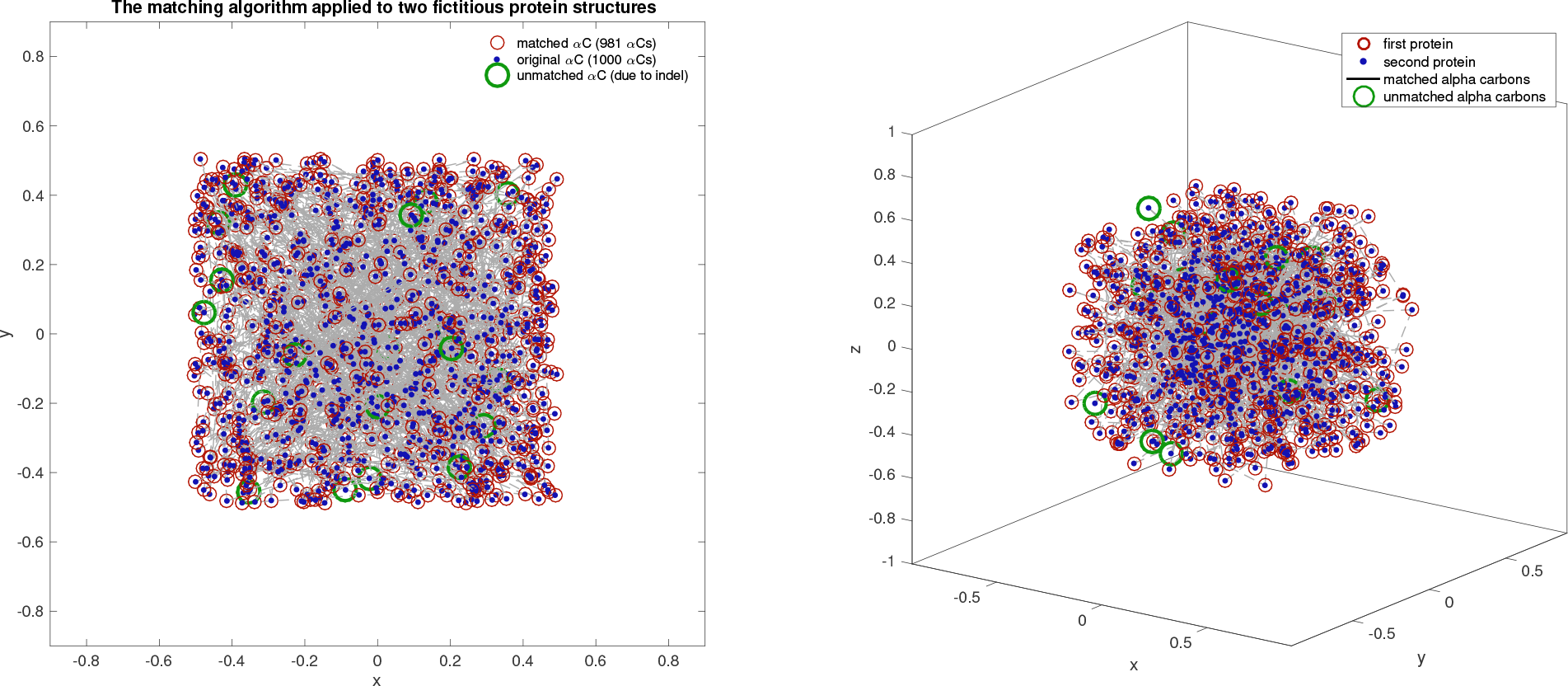
This is analogous to Figure S2 but for two structures of 1,000 and 980 randomly placed dots (a.k.a. vectors) in 3d. The algorithm correctly identifies 20 deleted dots in under 1s when implemented in Matlab 2016a running a MacBook Pro (Model MacBookPro12,1; Intel Core i5 2.9 GHz, 1 processor, 2 cores, L2 Cache per the Core 256k, L3 Cache 3 Mb, Memory 8 Gb RAM). The algorithm matches all but 20 missing 3d dots in two random datasets of 10,000 and 9,980 dots in under 30s on the same computer; the number of brute-force, vector-vector comparisons that would be needed to solve this problem naively amounts to over 4 × 10^61^. Depending on the mathematical metrics used, each of these comparisons would need a non-trivial vector operation like max over differences between sets of points of these sizes, or else that number of floating point multiplications if that metric were based on the Euclidean norm.

#### 1.4.1 Algorithm description

Given two vectorial datasets of arbitrary length, we would like to determine the best way of aligning the smaller dataset into the larger one such that that alignment is as close as possible. If the two datasets have the same length, then we simply seek the closest alignment which is a solved problem, as we discuss below. But in the former case their need not even be a unique solution to the problem as posed, so what follows represents a working heuristic to iterate towards a good alignment from some initial starting guess. Once this is done, we claim that alignment can be used to infer structural differences between the two datasets, for example in order to locate points that have been deleted or inserted, or simply moved within some localised region.

To perform these shape-based structural analyses, we need an algorithm capable of performing the following geometric matching problem. Given two un-ordered sets of vectors in **R**^3^, label them 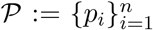 and 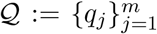, where *n* ≠ *m*. We need a readily-computable metric between 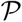 and 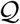 and for this we apply the following iterative heuristic to the rectangular Hungarian (munkres) algorithm.

By ‘un-ordered’ here we mean the subscripted index of either set is not used in the matching algorithm. Thus, if we change two indicies explicitly by applying a symmetry, *σ*, so that

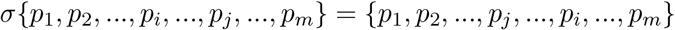

where *i* ≠ *j*, then an optimal assignment of 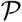 to 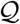 is based on geometric positions and not on the indices themselves. Thus, matching 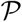 to 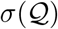 should give the same results as matching 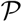 to 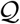. To paraphrase this, 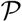 and 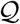 are considered to be unordered lists or sets here, not ordered vectors.

We use the Euclidean norm ‖ · ‖ defined on **R**^3^ and inherit a so-called Hausdorff distance from it that acts on sets, *d*, via

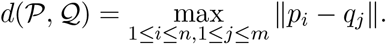

As a result, 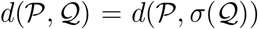 for any symmetry *σ* because the maximum in this definition is taken over all possible pairings between 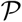 and 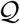.

We now seek a rigid-body transformation, *R*, that places 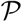 as close to some 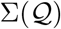 as is possible, where Σ = *σ*_1_ º σ_2_ º … º *σ*_*k*_ is some permutation of the indices. To achieve this we first respect the indexing and solve, for an unknown transformation *R*, the problem

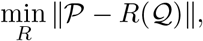

where the minimum is taken over all affine mappings, *R*(*v*) = *τ* + *Av*, that satisfy det(*A*) = 1. Here *τ* is a translation vector in **R**^3^ and *A* is a linear mapping of **R**^3^ to itself that preserves Euclidean distances, so *A* satisfies *A*^*T*^ *A* = *I*, and we apply *R* point-wise to 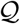, meaning 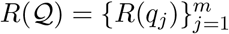.

This minimisation problem can be solved with a single singular-value decomposition (SVD) and the Matlab function matchPoints3d.m from the website https://github.com/rebear217/PhageMFiles does this. The assignment problem that optimally matches the labels of each element of 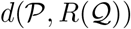 can then be solved with a single application of the Hungarian algorithm, which yields a candidate for Σ that matches labels {1, 2, 3, …, *m*} to some re-ordering of itself, as output, where Σ and *R* together achieve the minimum

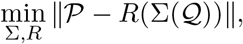

where Σ and *R* are as described above.

To try and solve this minimisation problem in general, even if *n* = *m*, and certainly if *n* ≠ *m*, the above algorithm must be extended to be iterative. For in the first case, *n* = *m*, there could be a better way of swapping two points with each other and reducing the distance between the resulting point sets, even without changing *R*. In the second case, *n* ≠ *m*, we then have a correspondence of our minimisation problem with the comparison of two proteins of different sizes that have insertion or deletion mutations. In this case, there is no way of knowing which indices in one protein can be matched with indices in the other.

So, to seek solutions to this with low computational cost, even if the algorithm is not guaranteed to converge theoretically, we apply the following heuristic. First, note that if *m* = *n* + 1 then there are *n* ways of choosing subsets of 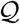, of size *m*, that have size *n*, each choice simply leaves out one of the points in 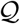; hence there are ^*n*^C_1_, or *n*, ways of doing this. If *m* = *n* + 2 then there are ^*n*^C_2_ = *n*(*n −* 1)/2 = *O*(*n*^2^) ways of choosing subsets of 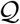 to match the size of 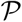 and, for general proteins of size *n* and *m* with *m* > *n*, there are *O*(*n*^*m*−*n*^) ways of doing this.

To overcome this combinatorial difficulty, the heuristic that iteratively compares subsets of 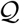 to 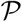, assuming the former is the larger of the two sets, is the following algorithm. It is based on the idea of starting with a random initial guess for the right number of points in the larger set that match the smaller set, which leaves some number of points ‘left aside’. We then update this each set sequentially by finding the optimal match of the current guess to the smaller set and ‘rejecting’ the right number of points that match the worst, replacing them with the set that is currently considered to be ‘left aside’. We call these the ‘keep for now’ (i.e. *K*_*j*_) and ‘leave aside’ (i.e. *L*_*j*_) sets in the following algorithmic pseudocode.

First let 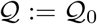, set *j* = 0, then

1. choose *n* random points from 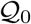 and call this set 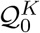 and call the remaining *m − n* points 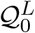; note^1^.
2. Let *K*_*j*_ and *L*_*j*_ denote the indicies of the two subsets of points, 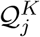 and 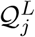 respectively.
3. Use SVD to determine the *R*_*j*_ that satisfies

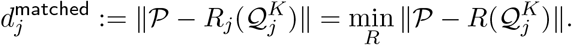
4. Now use *R*_*j*_ to shift 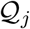 closer to 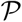 : set 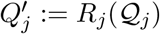 and define

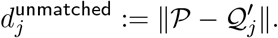
5. Let *K*_∗_ and *L*_∗_ be the sets of indicies, *K*_*J*_ and *L*_*J*_, which realises the shortest observed matching distance so far, meaning if we define the best distance observed so far

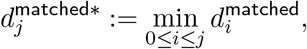

then 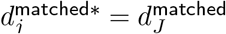 for some index *J* ≤ *j*; also set 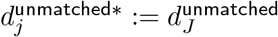
6. Now perform the rectangular Hungarian (munkres) algorithm that optimally matches each element of 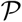 to one in 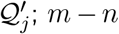 elements of the latter remain unmatched. The former set is *K*_*j*+1_ and the latter set is *L*_*j*+1_, 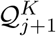 (matches) and 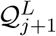 (non matches) are analogously defined by munkres.
7. If *K*_*_ = *K*_*j*+1_ and so *L*_*_ = *L*_*j*+1_ then terminate the algorithm with 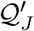 as the best match within 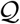 to the points in 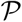; note^2^.
8. Otherwise, increase *j* by 1 and proceed to step 2.

At this point, we write the best observed distances, 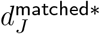 and 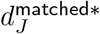, as *d*^matched^ and *d*^unmatched^.

While this algorithm is not proven to converge to an optimal solution, nor is an optimal solution to the matching problem sure to be unique (imagine trying to optimally match 3 of the vertices of a square back into the square), in practise the algorithm proceeds while 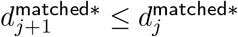 is satisfied and it is therefore a non-strict descent method within the space of possible matches.

Figure S2 gives an example of the algorithm applied to synthetic data in the shape of a figure ‘8’, showing that it can discriminate between matching sub-structures and unmatched ‘outliers’, shown as green circles. Figure S3 discusses what happen when applying the algorithm to matching problems with 1,000 and 10,000 points where it successfully 20 identified deleted 3-d points from datasets of each of those sizes with little difficulty in a matter of seconds, rather like finding a needle in a haystack.

To conclude, this algorithm improves vastly upon random matches between LamB protein structures with indels but we do not know if better matches might exist. (To visualise this, plots of structures, like those in Figure S10 and S11, show how closely the wild-type proteins match their mutants following a passage through the above algorithm.) After all, the algorithm can only converge to locally optimal solutions at best. However, at the heart of the matching algorithm is an iteration based on rigid transformations that are, by definition, non-expansive as they are norm preserving. It is therefore possible that geometric ‘genericity’ conditions on the datasets could be indentified whereby the matching problem is known to have a unique solution and where an algorithm like the one above determines it. However, this is beyond the scope of the present article but answers may lie in the theory of fixed points of non-expansive operators.

### Protein Hotspots

Now we have a way of matching two proteins, we can define a hotspots for each protein match: if 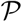 and 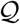 denote, respectively, a wild-type protein structure and a mutant structure, let *d* now be any well-defined measure of distance between the two proteins (i.e. a metric on point sets). The above matching algorithm defines measures *d*^matched^ and *d*^unmatched^ that could be used for *d* and in the article we use the former of these, even though it is not a distance metric in the strict sense (because it operates on sets of different sizes).

Now, if for some *α* ∈ (0, 1) there are points 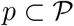 such that

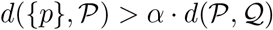

then each such *p* is said to be an *α*-hotspot. This condition formalises the idea that a fraction *α* of the entire distance from 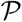 to 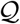 comes from just *p*’s contribution, making the latter a potentially larger distance from 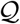 than ‘it should be’, justifying the term hotspot. Note if *α* is too large then there may be no hotspots at all and if *α* is too small then all points will be deemed hotspots; neither of these situations is desirable so we implemented the following heuristic.

The following rationale for the choice of hotspots is merely informal, it is not mathematically precise. If there are *n* points in each protein structure 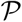 and 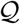 then a uniformly (i.e. randomly) distributed, between-points distance metric would have an expected distance that accounts for approximately 100 × 1/*n*% of the total distance per point. As the protein structures considered in this paper have *n* ≥ 400 then a value of *α* > 1/400 ≈ 0.0025 would be the expected, per-point distance following a protein match. To find extreme hotspots, we chose a value of *α* = 0.01 which is, therefore, 4× greater than expected based on a uniform distribution, and we call them 1% hotspots throughout.

## 2 Additional Cost-of-Resistance Tests

We tested whether growth rate, biomass yield (cells produced per maltotriose supplied) or lag time would shows differences between the zero susceptibility (i.e. pan-resistant) and positive phage susceptibility groups and found no clear evidence of differences between them with respect to rate or yield. We did observe an increase in lag times associated with pan resistance, but only for 3 out of 8 of the maltotriose concentrations tested (significant with respect to Bonferroni correction), see Figure S5.

## 3 Figures

**Figure S4:**
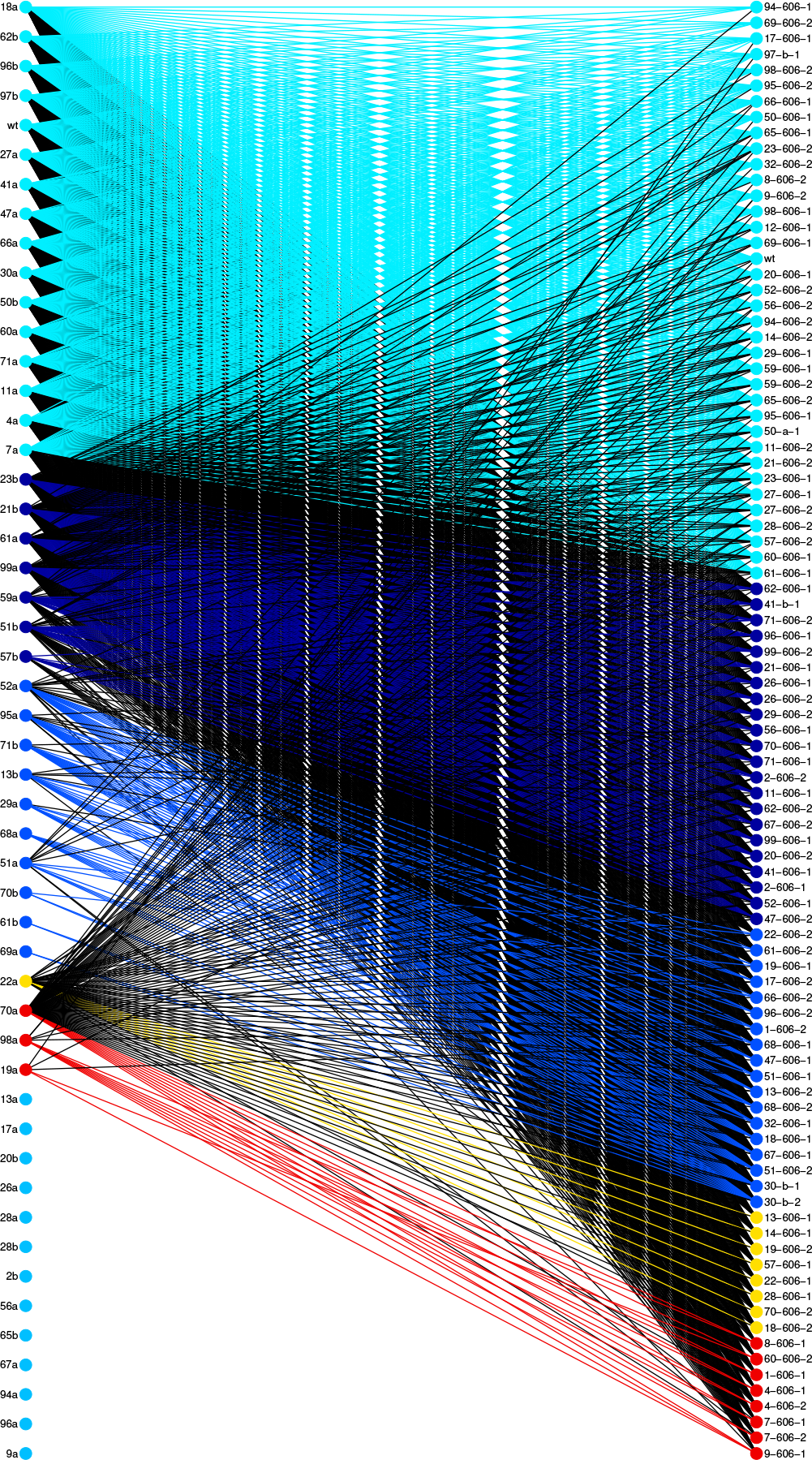
A bipartite graph representation of the infection matrix in Figure 2 based on the modularity analysis and associated colour coding illustrated in Figure 2(b). This graph shows library bacteria (left column) and phage (right) and a line indicates where a phage *can* infect a bacterium. Coloured lines denoted putative functional groupings into algorithmically-determined optimal sub-modules, *a.k.a.* potential locks and keys, whereas black lines denote possible infections outside those modules. The presence of so many black lines is consistent with there being no lock-and-key modular structure within the infection matrix because the latter is, in fact, nested. (So, a perfect lock and key system would have no black lines.)

**Figure S5:**
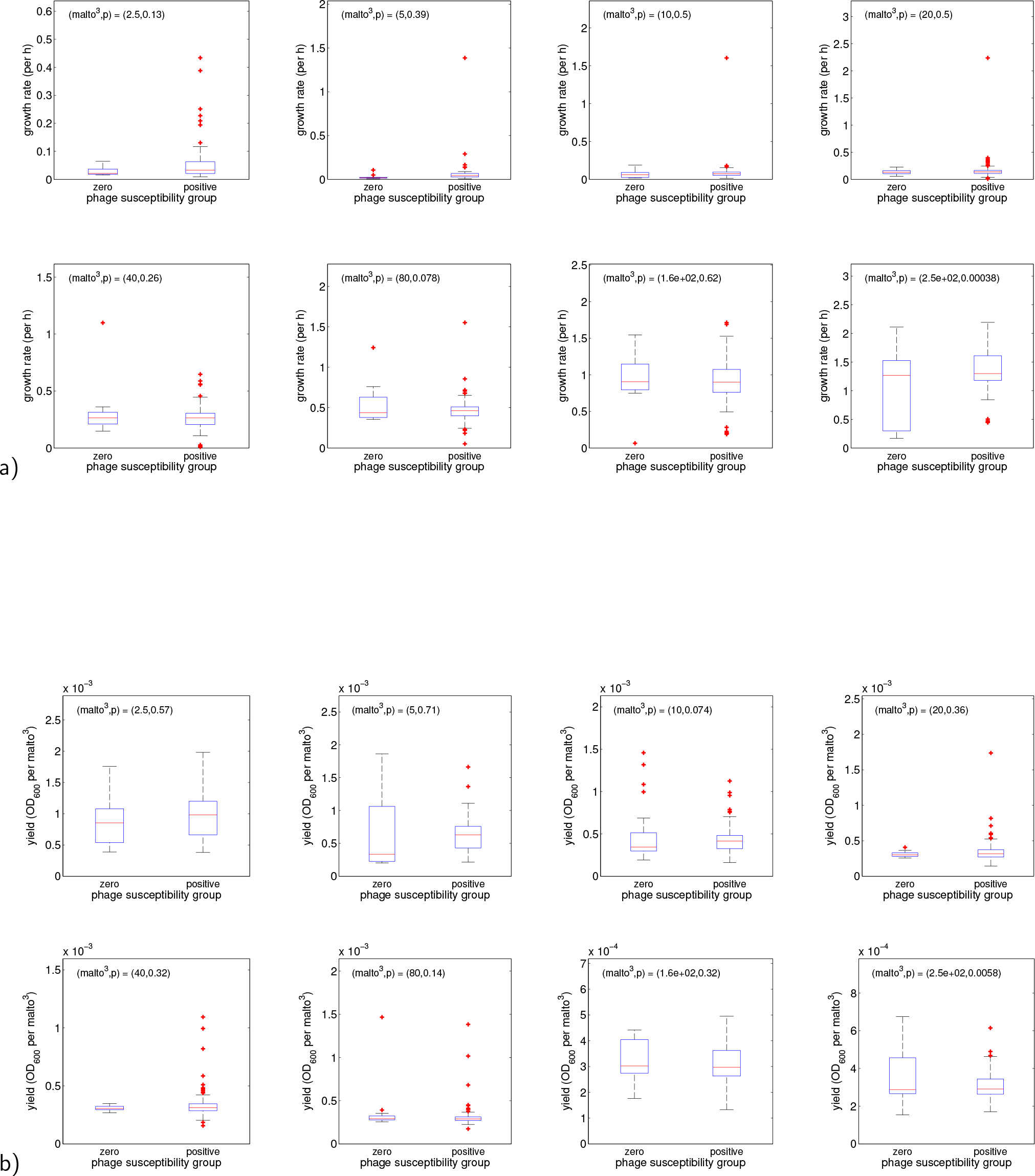
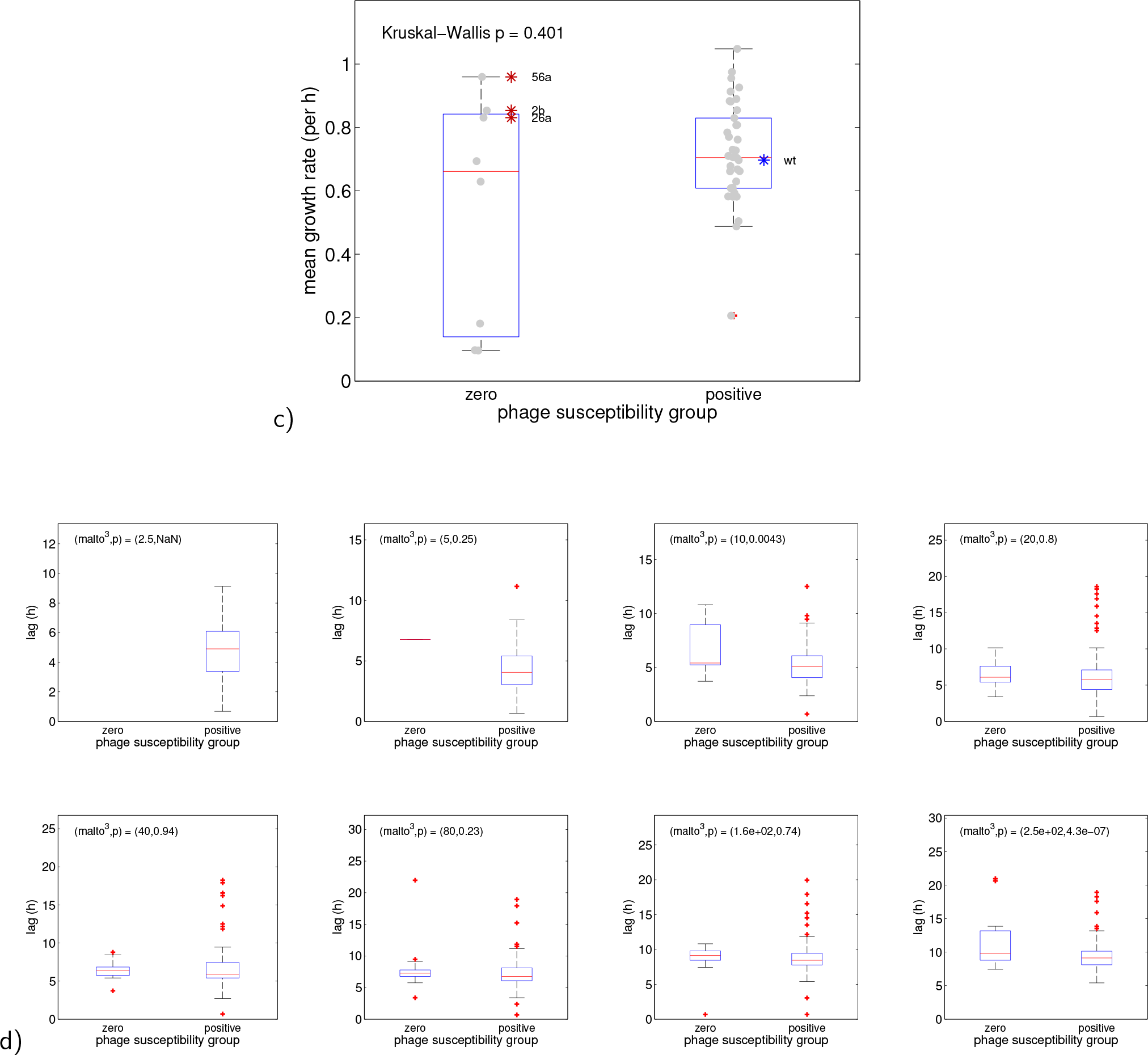
Pan-resistance has no growth rate nor yield costs, but does have a ~ 1h cost in lag. Bacterial growth rates (in (a)) and yields (in (b)) were determined in 8 different maltotriose concentrations that are quoted in each figure legend in *μg/ml*. Pan-phage resistant bacteria (zero group) were compared against phage-susceptible bacteria (positive group) to test for fitness costs of pan phage resistance but no evidence of these was found. c) We reach the same conclusions if (differences in) mean growth rate (i.e. averaged over all environments) is tested between zero and positive groups. d) A significant increase in lag time (~1h) is observed for the zero group in 2 maltotriose concentrations (namely 10 and 250*μg/ml*) and significant following Bonferroni correction.

**Figure S6:**
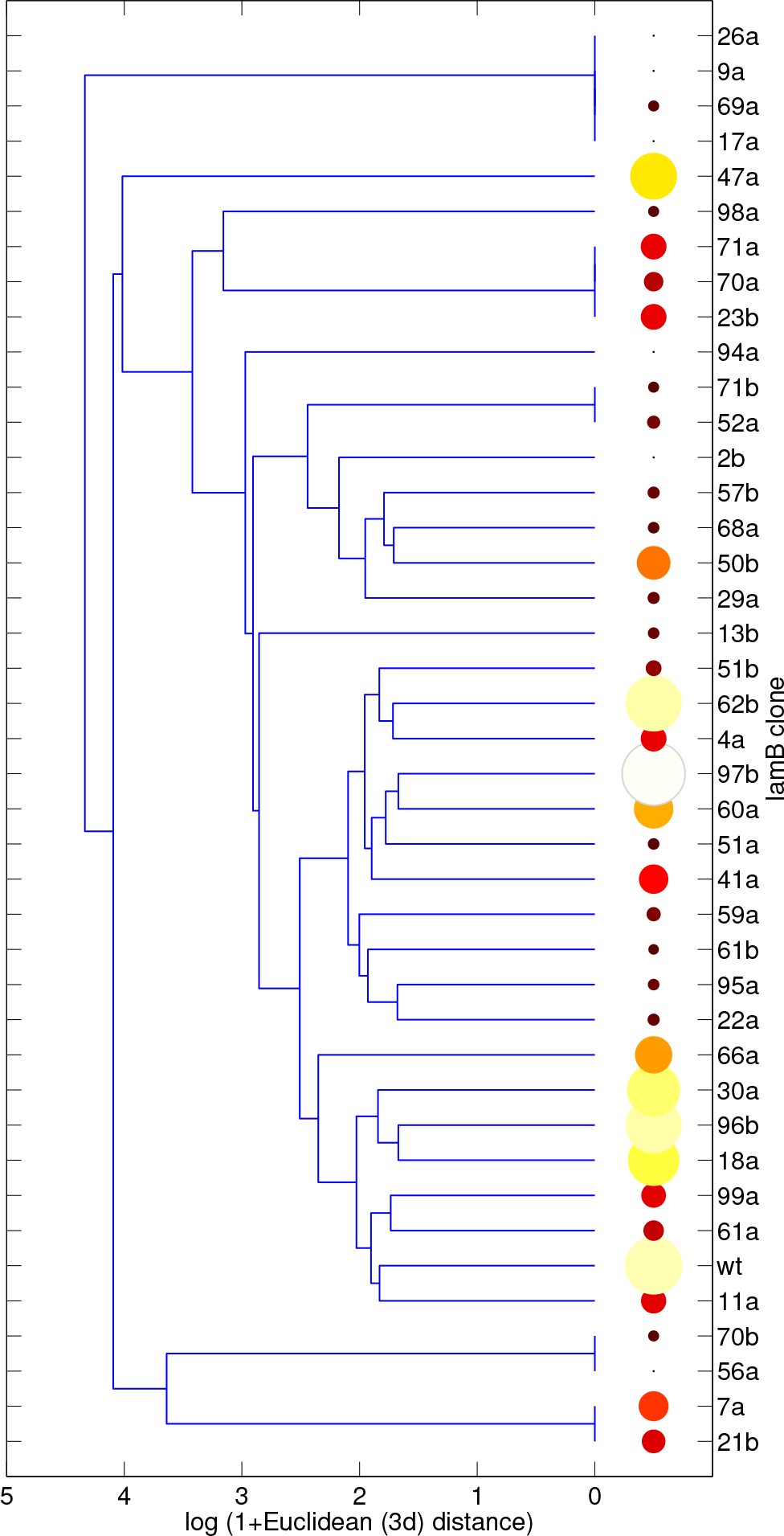
LamB protein distance-from-wild type correlates weakly with change in phage susceptibility. A dendrogram based on each bacterium’s LamB structure vector illustrates the weak correlation between structure and phage susceptibility. Each leaf has a size-coded dot where larger-whiter dots are more phage susceptible; the colour-code uses the black-red-yellow-white ‘hot’ colourmap whereby white is most susceptible. Note how the dots do not cluster according to like phenotypes around the dendrogram.

**Figure S7:**
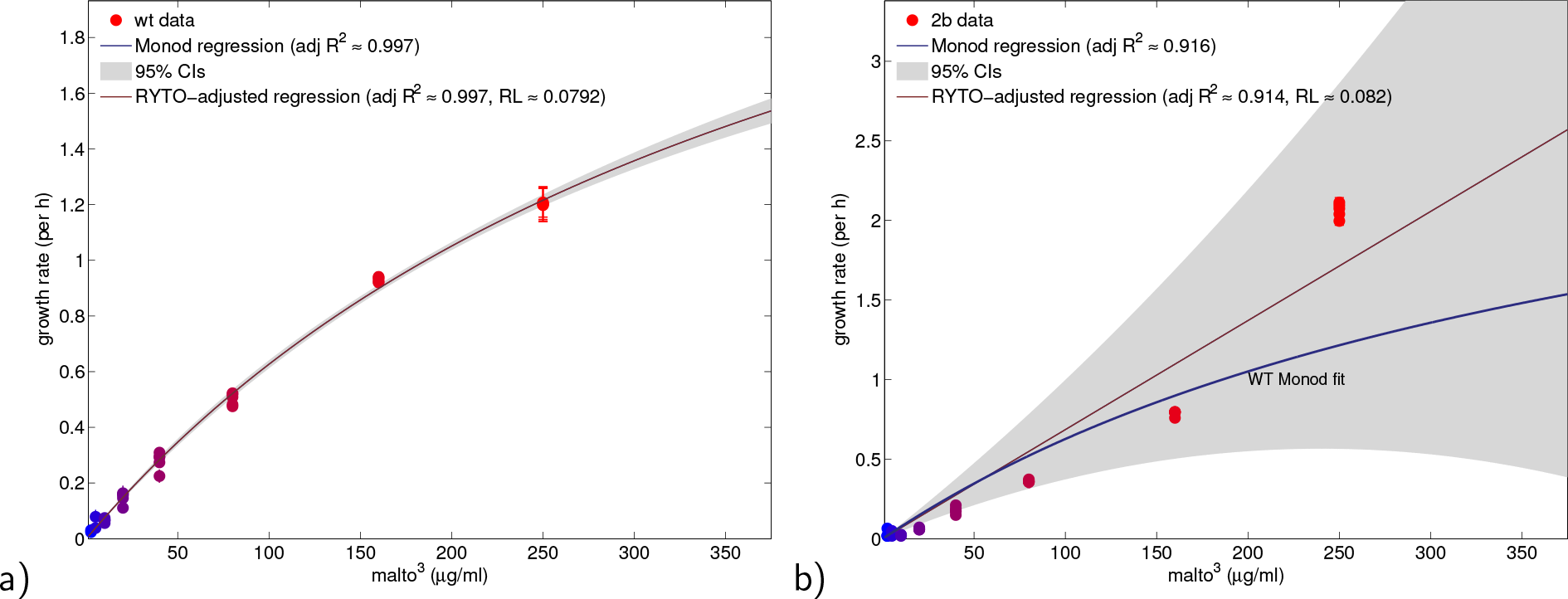
a) Growth rates versus maltotriose supply concentration for the wild-type library bacterium (REL606) measured in 8 maltotriose backgrounds. Growth rate data are dots, standard error is shown with *n* = 6 per condition). A claret line is a Monod regression to data and the grey area is a 95% CI of that regression. A second fit, called the RYTO-adjusted regression, is also fitted and shown in the image but this is visually almost indistinguishable from the first regression. Models for both regressions are described in the Methods (respectively, equations (5) and (7)). ‘RL’ in the legend is the ‘relative likelihood’ of the two models, the Monod regression being the more likely of the two, given these data. b) Growth rates of the pan-phage resistant bacterial genotype labelled ‘2b’ are shown here: this strain exhibits faster growth than the WT (see blue line) but at only one of the maltotriose concentrations tested, at all others it exhibits slower growth than the WT. (Note: the Monod and RYTO-adjusted regressions are relatively poor descriptors of the growth rate data here as they are not concave with respect to maltotriose concentration. Moreover, 250*μg/ml* does not appear to be close to the concentration at which growth rate is limiting for 2b, in contrast to the WT whose growth rate data are well-described by Monod regression.

**Figure S8:**
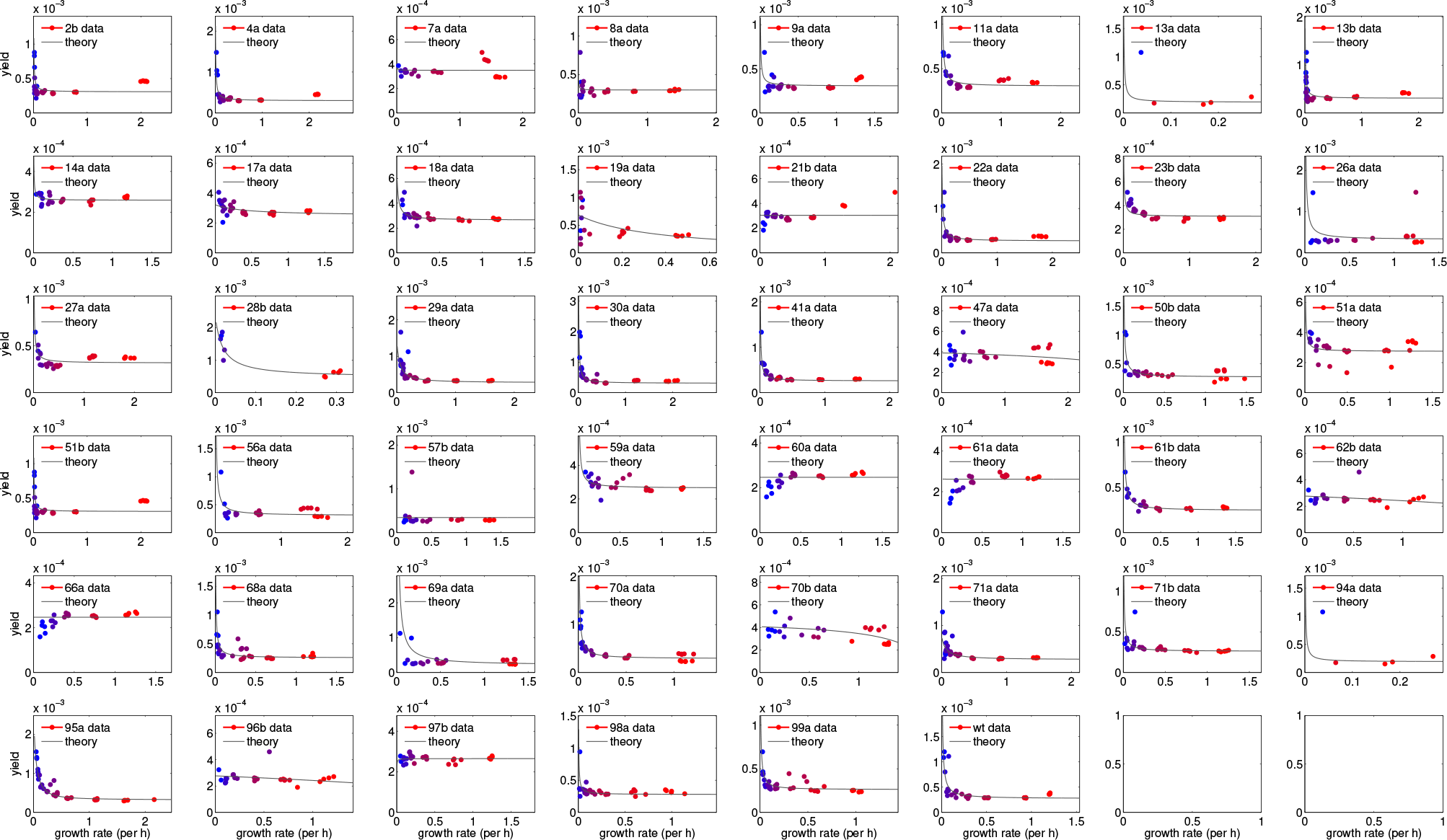
Rate-yield relationships for 46 bacterial library strains. Most datasets form rate-yields tradeoffs (RYTOs) where yield decreases with increasing growth rate that are well-described by trade off theory (equation (6)), but this is not true for all strains. Exceptions are strains for which yield increases along with rate and there are 3 of these, 60a, 61a and 66a. Note, yield is measured here in units of optical density (OD_600*nm*_) per *μg* of maltotriose and growth rate is measured per h.

**Figure S9:**
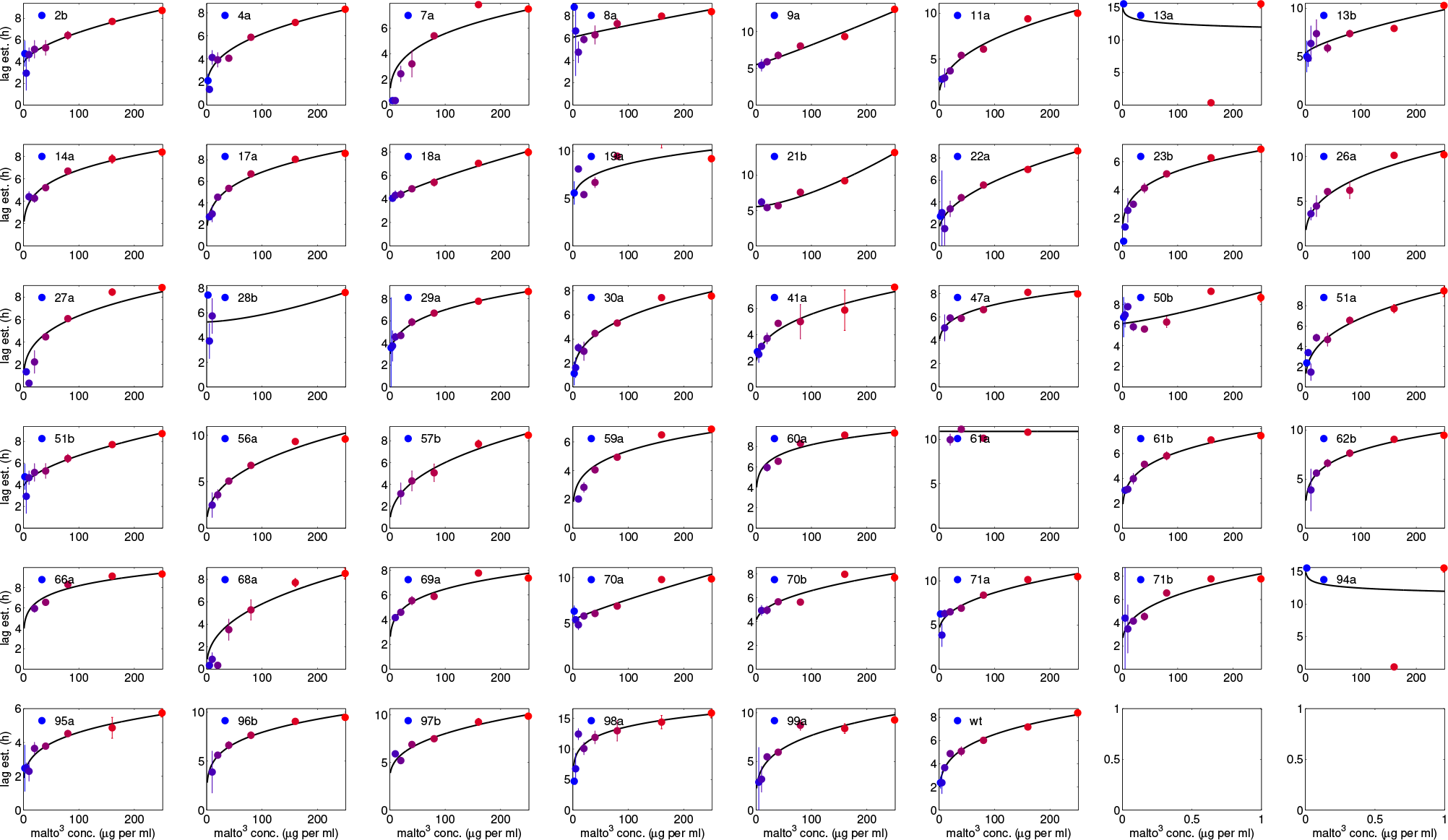
The lag-maltotriose supply relationships shown here of 46 bacterial strains in the library are presented for completeness. Note that growth rate increases with increasing maltotriose, whereas yield decreases, creating within-strain rate-yield tradeoffs (see Figure S8). As a result of that observation and the data here showing lag increases with maltotriose supply, lag will also correlate negatively with yield, but those data are not shown.

**Figure S10:**
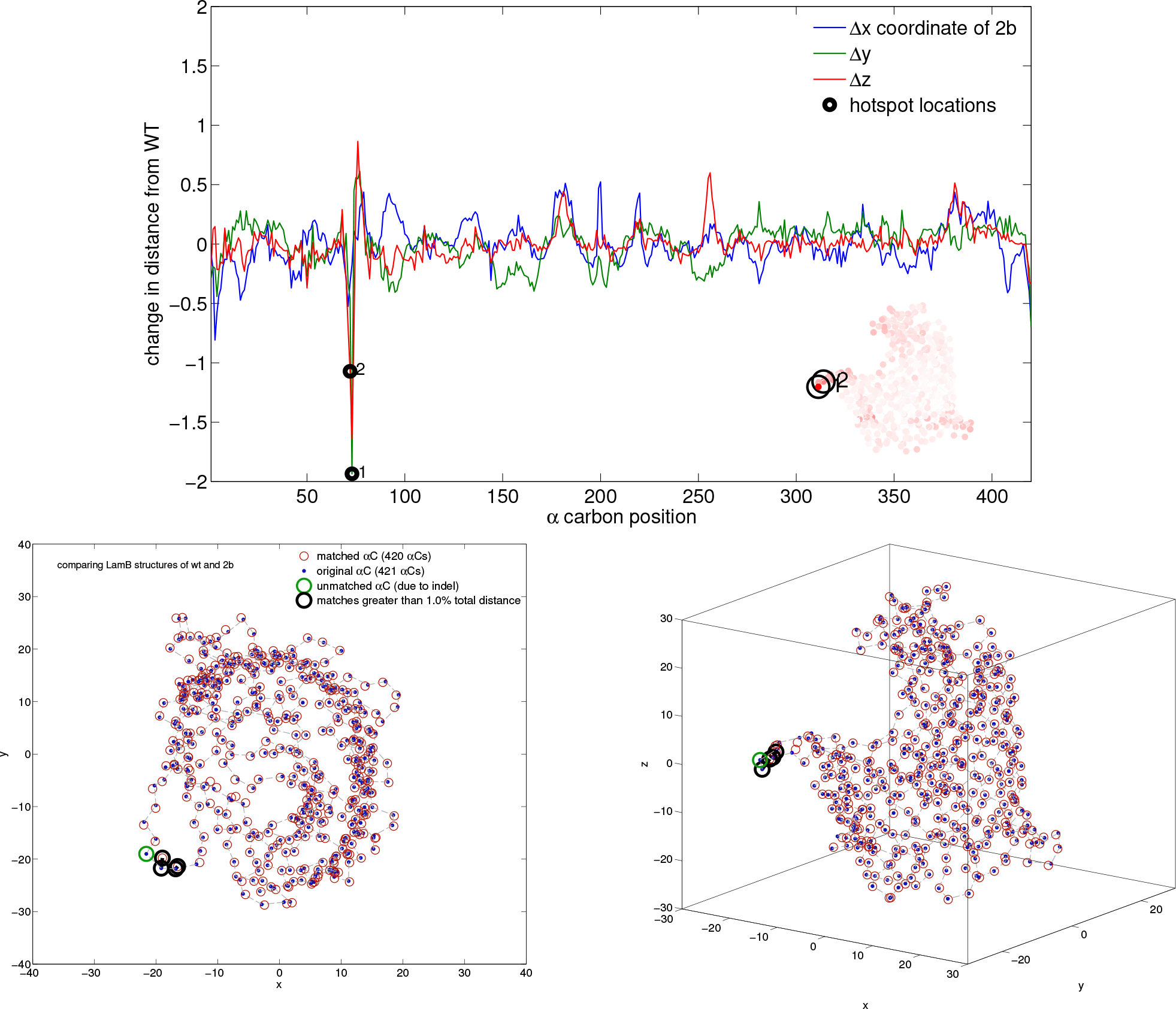
The analogous figure to Figure 5 from the main text but for bacterial strain 2b which, like 26a in that figure, has a localised change in LamB with respect to the wild-type ancestral structure.

**Figure S11:**
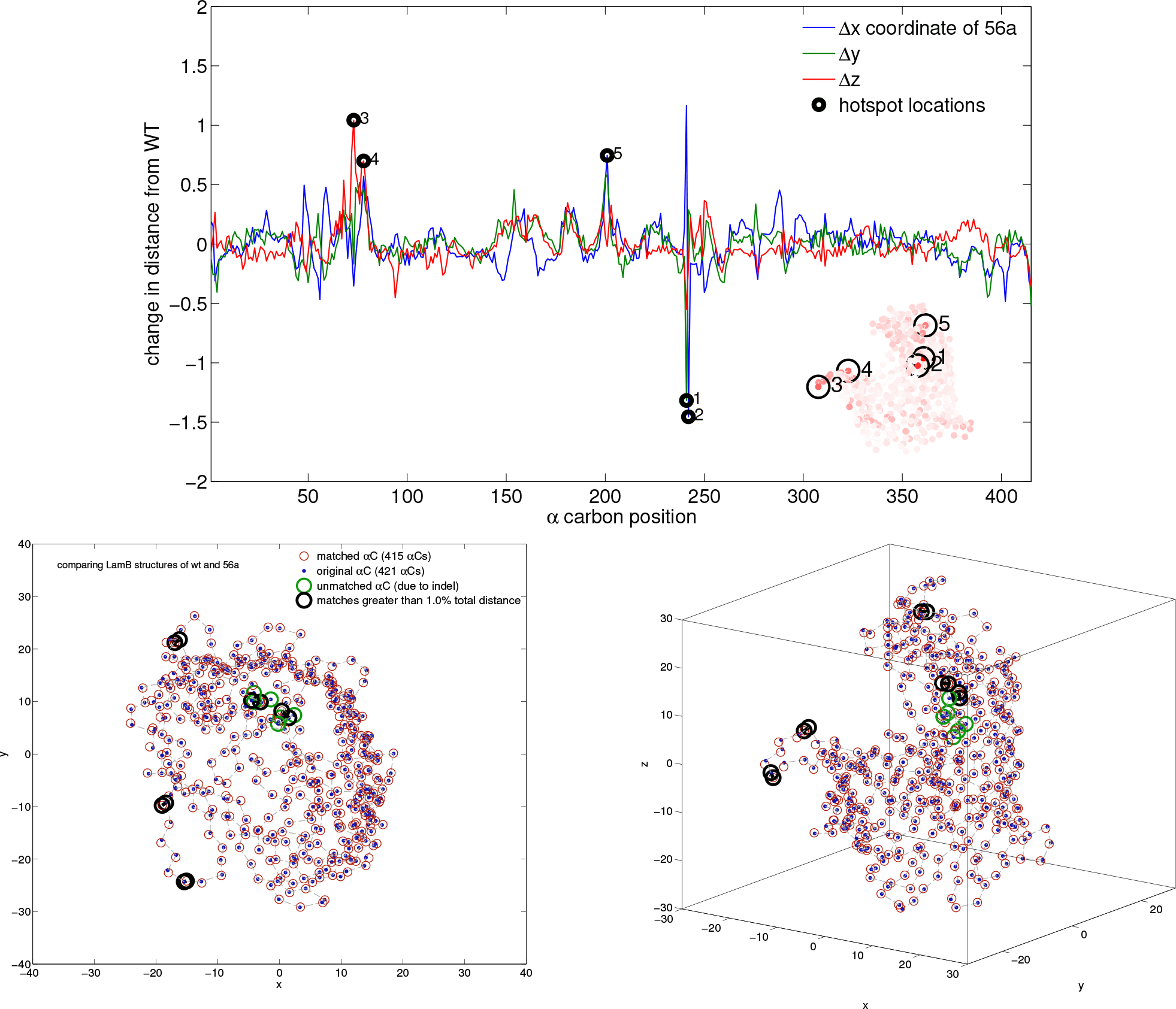
The analogy of Figure 5 from the main text but for bacterial strain 56a. The change in LamB with respect to the wild-type structure in this case is not as localised for 56a as it is for strains 2b (Figure S10) and 26a (Figure 5) although those changes on LamB arise at similar locations in the protein.

**Figure S12:**
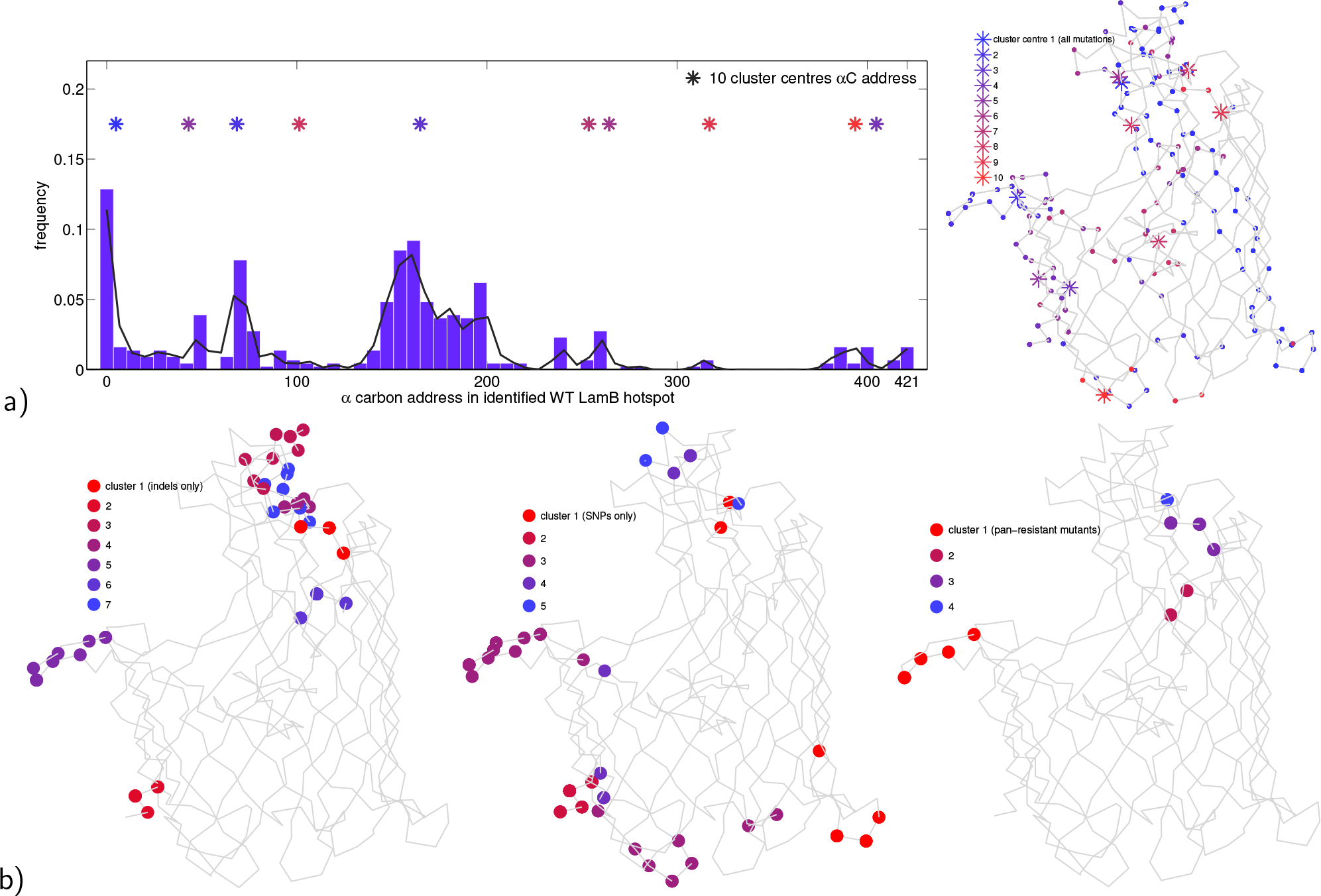
Different sub-groupings of the bacterial library have LamB changes that spatially cluster. a) Local maxima of the kernel density estimate (KDE) of the distribution of frequencies for which each LamB alpha carbon position appears in hotspots for the *entire* bacterial library (left) were used to conservatively (not optimally) identify as many alpha carbons as possible that could form putative LamB hotspots of large structural change. *k*-means clustering uses these locally maximal frequencies in the KDE to determine 10 hotspot cluster centres associated with those geometric changes (these are the coloured asterisks) and, as shown in the rightmost LamB structural plot, these appear in the greasy slide and outer loop regions. b) Re-performing the analysis for a) subsets of the bacterial library yield clusters of LamB hotpots associated with (left) indels, (middle) SNPs and (right) pan-phage resistance. The latter identifies changes in two extracellular regions of LamB that correlate with pan resistance; this may be due to mutations in these regions preventing phage from binding to the cell surface.

**Figure S13:**
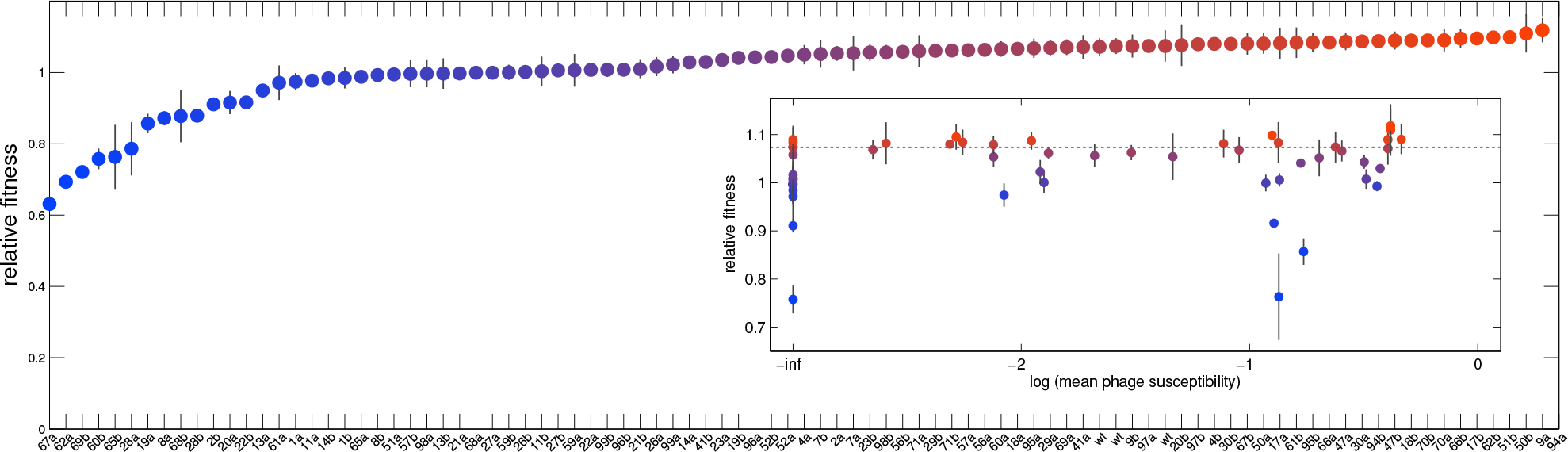
There is no correlation between bacterial competitive ability against the wild-type and phage resistance. All strains’ relative fitnesses in comparison to the WT are shown and plotted versus phage susceptibility. Relative fitness values are shown in the main plot, the inset shows the relationship between each relative fitness and that strain’s mean phage susceptibility against all phage; we note the absence of correlation.

**N.B.**: The following 3 figure pages contain all the geometric comparisons made between wild-type and mutant LamB structures. We present these here in order to demonstrate that the geometric matching algorithm has faithfully and automatically determined appropriate structural matches and not mis-matches. That this is true can be verified visually by noting that the two LamB geometries (one shown as blue dots and the other shown as red circles) are close to each other at all alpha Carbon locations for all mutants, and the black and green regions on each protein represent algorithm-determined outliers within those structures that do not match as well.

We present these for the reason that there are very many cases of toy exemplar matching problems that, when presented to the algorithm, it does not converge. This is not surprising and it happens because, for any two arbitrary point sets in 3d-space, the problem of finding the 2-point pairings between those sets which exhibits the minimal distance between those two sets is a non-convex optimisation problem that may not have a unique solution. The algorithm is not therefore guaranteed to converge and, when it does converge, it is not sure to find an optimal solution. It is therefore only by post-hoc validation that we can assess whether the algorithms have returned solutions appropriate to the problem, which the following figures do show.

**Figure S14:**
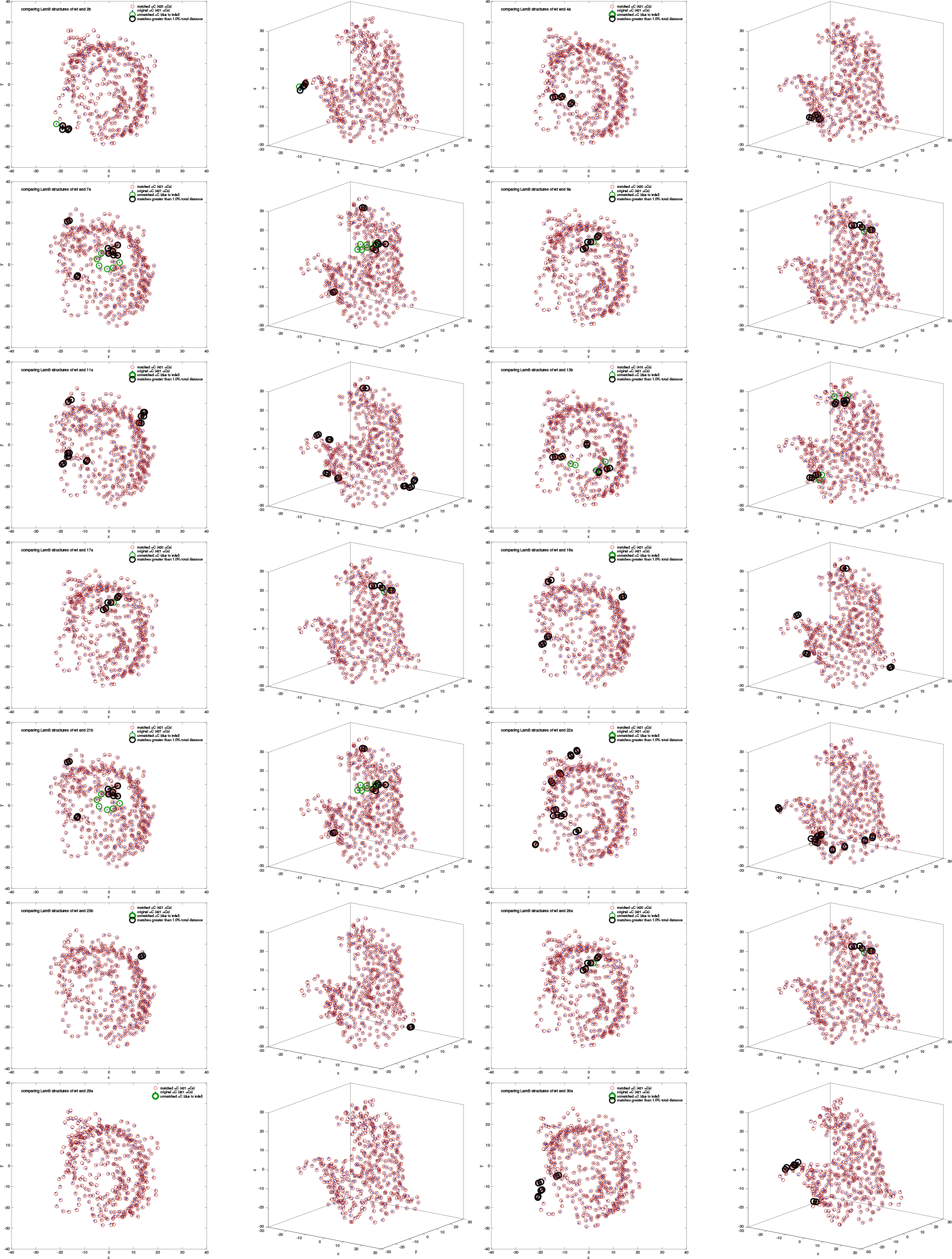
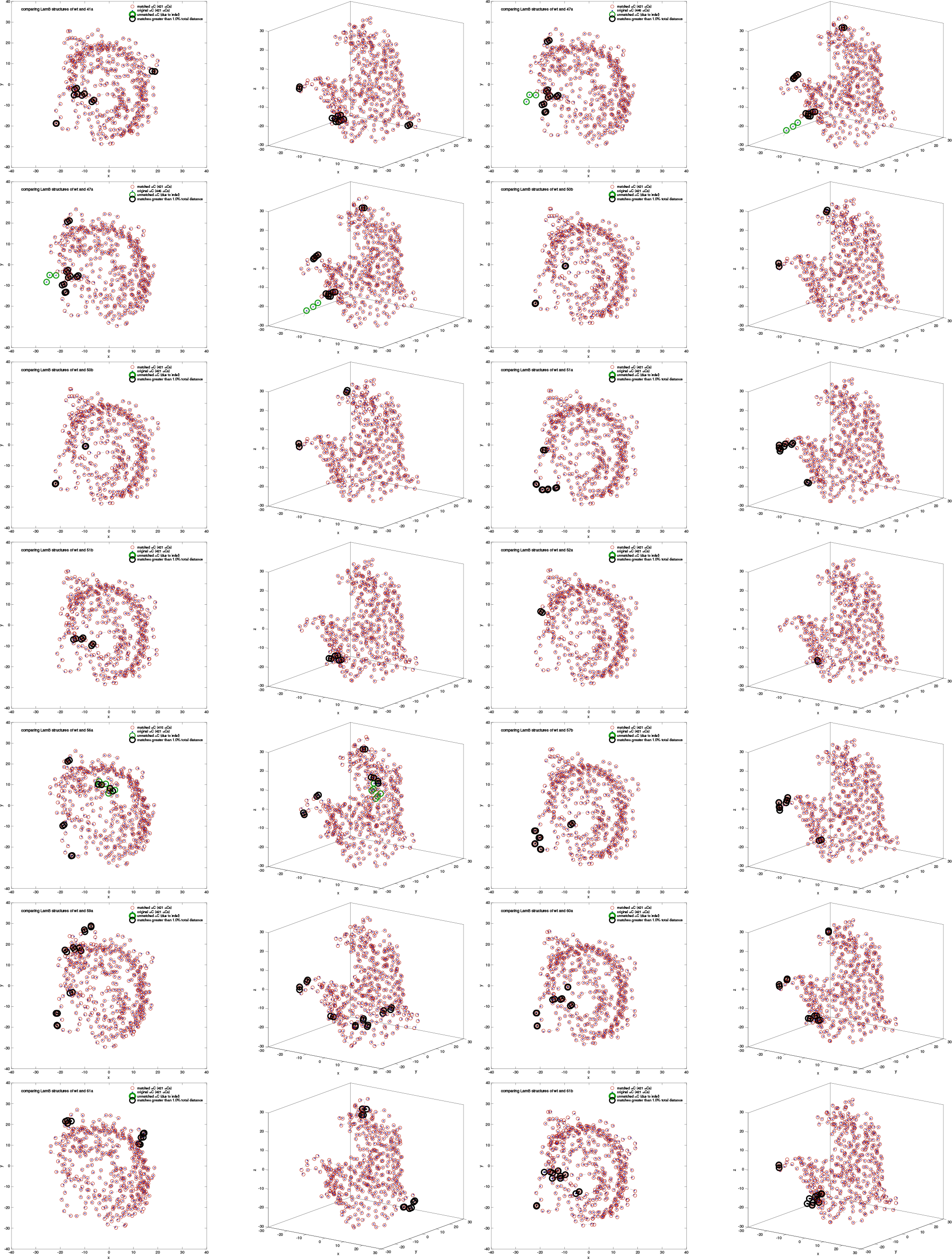
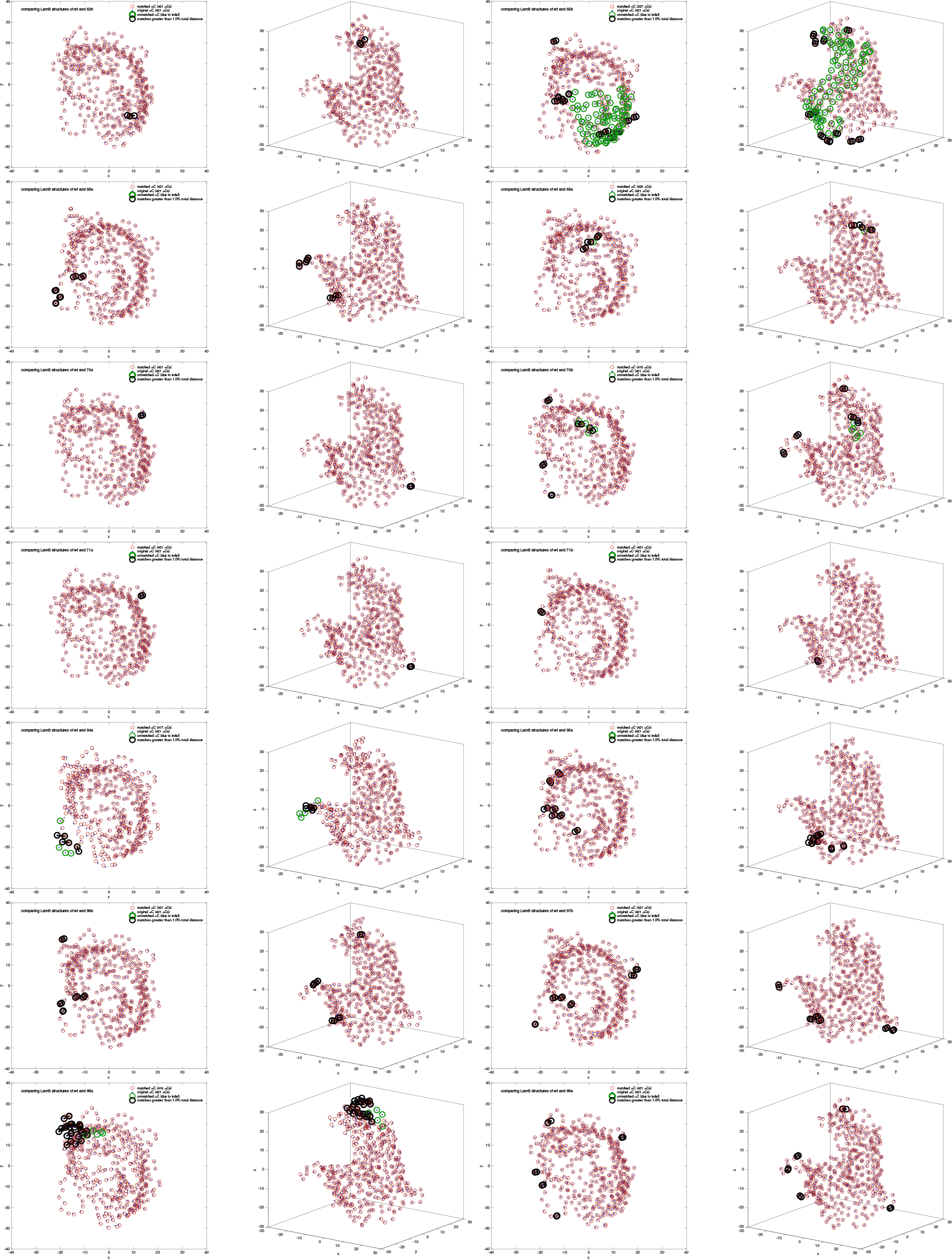
Forty-two wildtype-to-mutant LamB structural comparisons as determined by our matching algorithm.

## 4 Supplementary Tables

**Supplementary Table S1:** *E.coli* B(REL606) mutants are presented, the table showing their library identifier and mutations found in the genome.

| Strain | genome position | mutation | annotated | gene |
| --- | --- | --- | --- | --- |
| 2b | 4,227,194 | $\Delta 3$ bp | coding (292-294/1341 nt) | *lamB* |
| 3a | 1,882,795 | $\Delta 1,490$ bp | | *[manZ]–[yebN]* |
| 4a | 3,148,678 | G→T | Q211K | *yqjH* |
| | 3,894,997 | $\Delta 547$ bp | IS150-mediated | *[rbsD]–[rbsD]* |
|  | 4,227,756 | C→A | A285D | *lamB* |
| 8b | 1,882,827 | $\Delta 10$ bp | coding (363-372/861 nt) | *manZ* |
| | 3,894,997 | $\Delta 5,951$ bp | IS150-mediated | *rbsD–[rbsR]* |
| 9a | 3,894,997 | $\Delta 4,726$ bp | IS150-mediated | *rbsD–[rbsK]* |
| | 4,227,766 | $\Delta 3$ bp | coding (864-866/1341 nt) | *lamB* |
| 10a | 787,879 | $\Delta 12,090$ bp | | *ECB_00726–*<br>*ECB_00739* |
|  | 1,865,323 | A→T | S301C | *yeaX* |
|  | 1,883,048 | IS186 (–)<br>+7 bp | coding (584-590/861 nt) | *manZ* |
|  | 2,788,553 | T→C | D74G | *ygcW* |
| 15a | 1,882,844 | G→A | W127* | *manZ* |
| 16a | 3,482,762 | G→C | A360P | *malT* |
| | 3,894,997 | $\Delta 4,189$ bp | IS150-mediated | *rbsD–[rbsK]* |
| 19a | 3,894,997 | $\Delta 3,873$ bp | IS150-mediated | *rbsD–[rbsB]* |
|  | 4,227,513 | +T | coding (611/1341 nt) | *lamB* |
| 20a | 1,882,080 | +AC | coding (420/801 nt) | *manY* |
| 22b | 1,881,938 | $\Delta 1$ bp | coding (278/801 nt) | *manY* |
| 24a | 1,881,767 | +G | coding (107/801 nt) | *manY* |
| | 3,894,997 | $\Delta 154$ bp | IS150-mediated | *insK-5/rbsD* |
| 25a | 1,879,367 | $\Delta 3,809$ bp | | *[yoaE]–[manZ]* |
| 44a | 1,882,112 | $\Delta 1$ bp | coding (452/801 nt) | *manY* |
| 49a | 1,881,802 | +A | coding (142/801 nt) | *manY* |
| | 3,894,997 | $\Delta 3,166$ bp | IS150-mediated | *rbsD–[rbsB]* |
| 51a | 4,227,726 | C→A | S275Y | *lamB* |
| 52a | 904,341 | C→T | intergenic (+45/-13) | *ECB_00850/ECB_00851* |
|  | 2,103,918 | +CCAG | coding (185/216 nt) | *ECB_01992* |
| | 3,894,997 | $\Delta 4,631$ bp | IS150-mediated | *rbsD*–[*rbsK*] |
|  | 4,207,476 | C→T | A272V | *yjbB* |
|  | 4,227,711 | G→T | G270V | *lamB* |
| **54a** | 1,881,936 | $\Delta 11$ bp | coding (276-286/801 nt) | *manY* |
| | 3,894,997 | $\Delta 6,427$ bp | IS150-mediated | *rbsD*–[*yieO*] |
| **55a** | 787,879 | $\Delta 12,090$ bp | | *ECB_00726*–<br>*ECB_00739* |
| **56a** | 1,630,681 | IS1 (+) +9 bp | coding (422-430/1020 nt) | *rspB* |
| | 4,227,698 | $\Delta 18$ bp | coding (796-813/1341 nt) | *lamB* |
| **60b** | 3,483,087 | $\Delta 2$ bp | coding (1403-1404/2706 nt) | *malT* |
| | 3,894,997 | $\Delta 4,382$ bp | IS150-mediated | *rbsD*–[*rbsK*] |
| **61a** | 4,227,420 | A→C | E173A | *lamB* |
| **61b** | 3,894,997 | $\Delta 1,345$ bp | IS150-mediated | *rbsD*–[*rbsA*] |
|  | 4,227,461 | G→T | D187Y | *lamB* |
| **65b** | 517,069 | IS1 (+) +9 bp | coding (382-390/1236 nt) | *allC* |
|  | 2,745,465 | A→G | intergenic (-170/-42) | *hycA/hypA* |
| | 3,894,997 | $\Delta 98$ bp | IS150-mediated | *insK-5/rbsD* |
|  | 4,227,961 | C→G | Y353* | *lamB* |
| **67a** | 3,894,997 | $\Delta 6,815$ bp | IS150-mediated | *rbsD*–[*yieO*] |
|  | 4,227,038 | C→T | Q46* | *lamB* |
| **68a** | 4,227,717 | C→T | S272L | *lamB* |
| **89a** | 3,894,997 | $\Delta 282$ bp | IS150-mediated | [*rbsD*]–[*rbsD*] |
| **98a** | 3,894,997 | $\Delta 3,338$ bp | IS150-mediated | *rbsD*–[*rbsB*] |
| | 4,227,605 | $\Delta 14$ bp | coding (703-716/1341 nt) | *lamB* |

**Supplementary Table S2:**
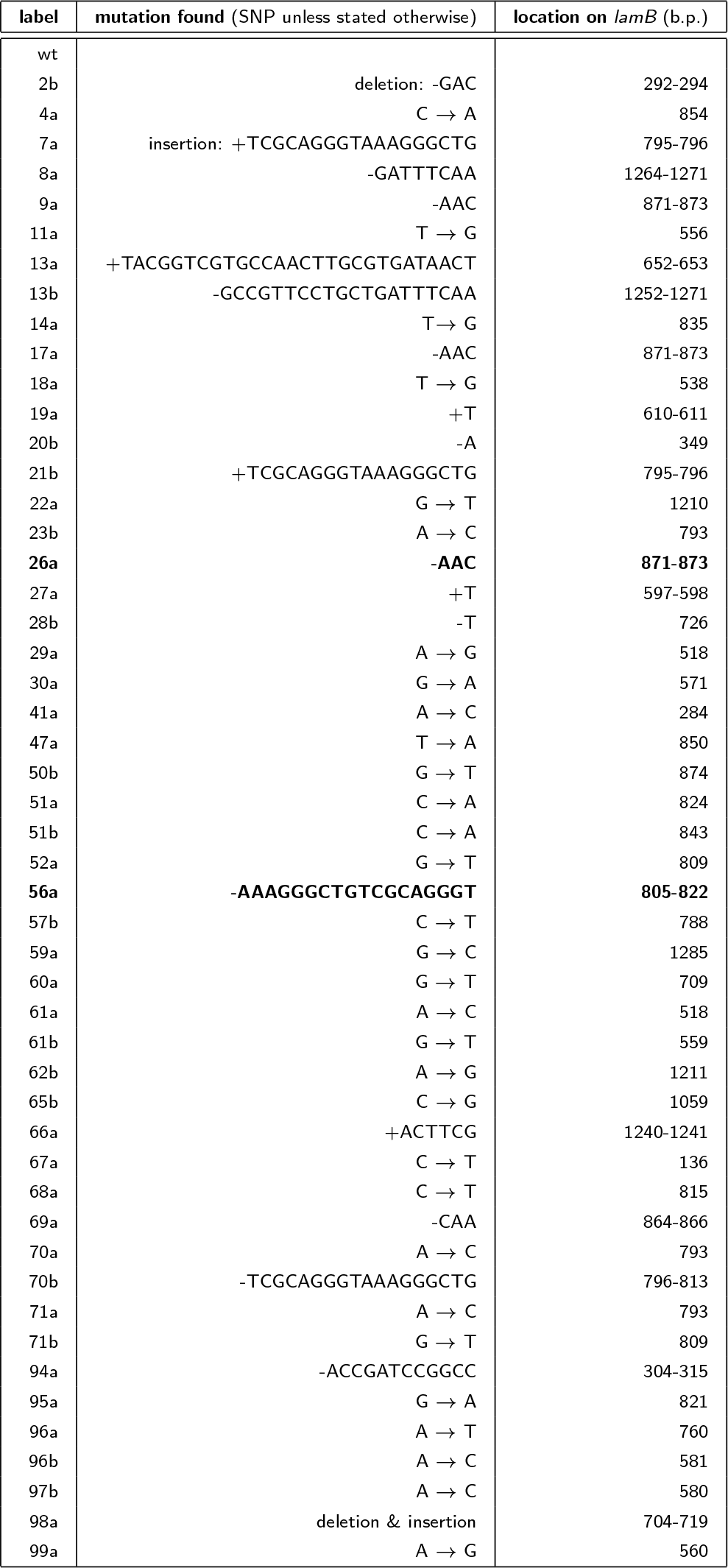
*E.coli* B(REL606) mutants showing their library identifier and mutations in *lamB*. The ‘Darwinian Monster’ phenotypes from the main text are highlighted using bold font.

## 5 Fitness can increase from an uptake-reducing transporter mutation in the presence of a rate-yield tradeoff

The main text asks that the change in growth rate that results from a small change in maximal uptake rate, *dV*, can be approximated by calculus, whereby:

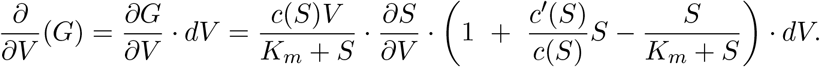

We wish to know whether growth rate, *G*, can go up as *V* goes down: can 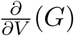 be positive when uptake rate decreases (itself postulated to be the result of a phage resistance mutation that impairs transporter function) and therefore *dV* < 0? For this to occur, because *V* > 0, *S* > 0, *K*_*m*_ > 0, *c* > 0, 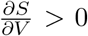, we need the expression

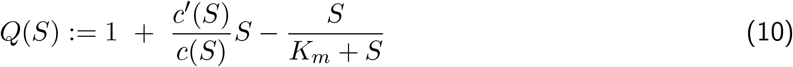

to be negative where *c* is defined in equation (6) of the main text: *c*(*S*) = (*c*_hi_ + *c*_lo_*pS*)/(1 + *pS*).

So, first do a little factoring and write

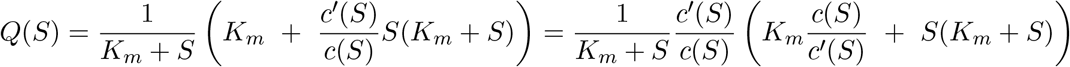

and spot that 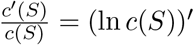, where (as is standard) a dash denotes derivative. Now

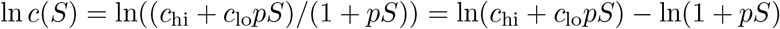

which yields

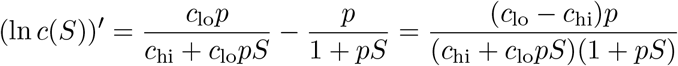

whence

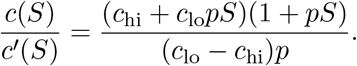

For the RYTO to apply, we require the value of *c*(*S*) to be higher for lower values of sugar, *S*, which is equivalent to *c*_hi_ *> c*_lo_, so we now assume this. Given that *p* ≥ 0, and so *c* is a decreasing function, namely *c*^*t*^(*S*) < 0, we have

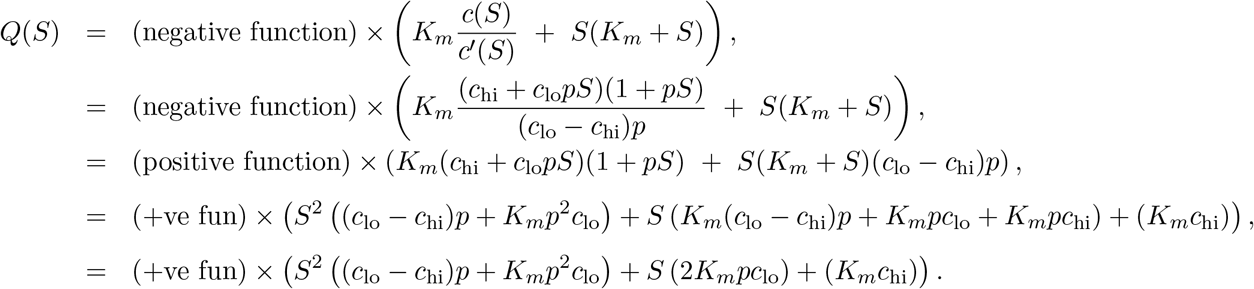

If we define the quadratic expression

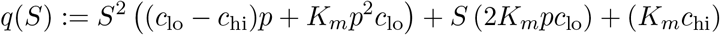

then *q* and *Q* have the same sign. Moreover, *q*(0) = *K_m_c*_hi_ > 0 and *q*^*t*^(0) = 2*K_m_pc*_lo_ > 0, which means that *q*(*S*) is positive and increasing when *S* = 0 which, given that it is a quadratic function, also means that it can only become negative if it has a maximum value for *S* > 0. In other words, the solution of *q*^*t*^(*S*) = 0 must occur for *S* > 0. A short calculation shows this requires

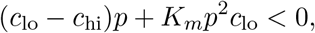

or

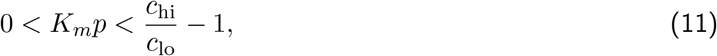

and in this case *q*(*S*) < 0 provided *S* > *S*_0_ where *S*_0_ is the solution of *q*(*S*_0_) = 0.

Now, Figure S8 shows in data that *c*_hi_ is about thrice the value of *c*_lo_ for the wild-type strain which gives an empirically-derived inequality of

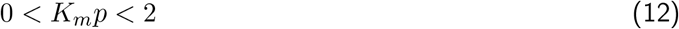

for which mutations in a maltotriose transporter of small negative effect, and which increase phage resistance, can also increase bacterial fitness but *only* provided maltotriose concentration is large enough.

In the situation where condition (11) fails, for instance *c*_hi_ = *c*_lo_ where there is no rate-yield tradeoff, then there would be no region of sugar concentrations for which a mutation with *dV* < 0 can increase growth rates.

## 6 More tradeoffs cause more tradeups

Suppose that traits *X* and *Y* are traded, so that *X* = *f* (*Y*) where *f* (*·*) is a one-dimensional trait relationship that is decreasing: *f* (*Y*_1_) *> f* (*Y*_2_) whenever *Y*_1_ < *Y*_2_ or, if *f* is a smooth function, it has negative derivative *df* / *dY*. Now suppose that traits *Y* and *Z* are also traded: *Y* = *g*(*Z*) where *g* is a decreasing function so that *dg/dZ* is negative. This places a constraint on how *X* and *Z* must interact as traits: *X* = *f* (*g*(*Z*)) and calculus says that

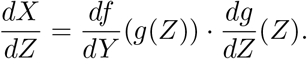

Inspecting the signs of these derivatives shows that *dX/dZ* is positive because it is the product of two negative quantities, thus *X* and *Z* engage in a ‘tradeup’. For this reason, tradeoff relationships are not transitive between trait pairs, however tradeups are.

This observation shows that if multiple traits interact (call them *A, B, C* and *D*, see Table S3) and all of *B, C* and *D* trade off with trait *A*, then the trait correlations between *B, C* and *D* must all be positive in order to meet the above transitivity properties which are the result of pairwise parity relationships that must hold pairwise between traits. Thus, wherever tradeoffs are common between traits, tradeups should be equally common.

**Supplementary Table S3:**
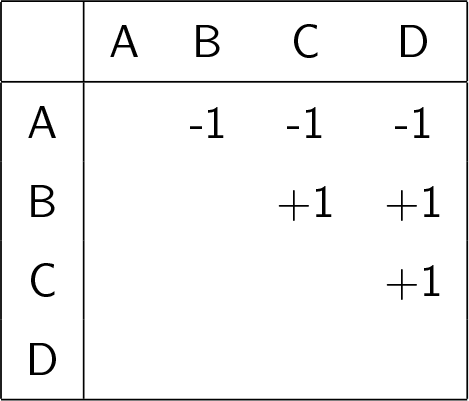
‘Parity relationships’ in a 4-trait system showing tradeoffs (−1) and tradeups (+1) between each trait, denoted *A, B, C* and *D* that interact pairwise.

It is exactly this argument in the main text by which we argue that mutations providing resistance to phage can interact with other traits, and their respective tradeoffs, and so conspire to create downstream growth rate benefits, as Table S4 illustrates. In it, we assume (see the red coloured −1) that sugar uptake rate goes down as per-phage resistance increases (by mutation) due to a structural change in the protein that is, we assume, *de facto* unlikely to confer the dual benefits of increased resistance and faster sugar uptake. Indeed, such mutations that directly bestow dual benefits, if possible, could provide a further, and simpler, explanation of some of our data.

To re-iterate, the need for two tables arises because growth rate is a composite trait of uptake rate and cell yield which are themselves both tradeoffs and it is the relative strength of the latter two, which changes with resource concentration, which dictates the nature of the growth rate tradeoff. Indeed, the latter can become a tradeup under some circumstances.

**Supplementary Table S4:**
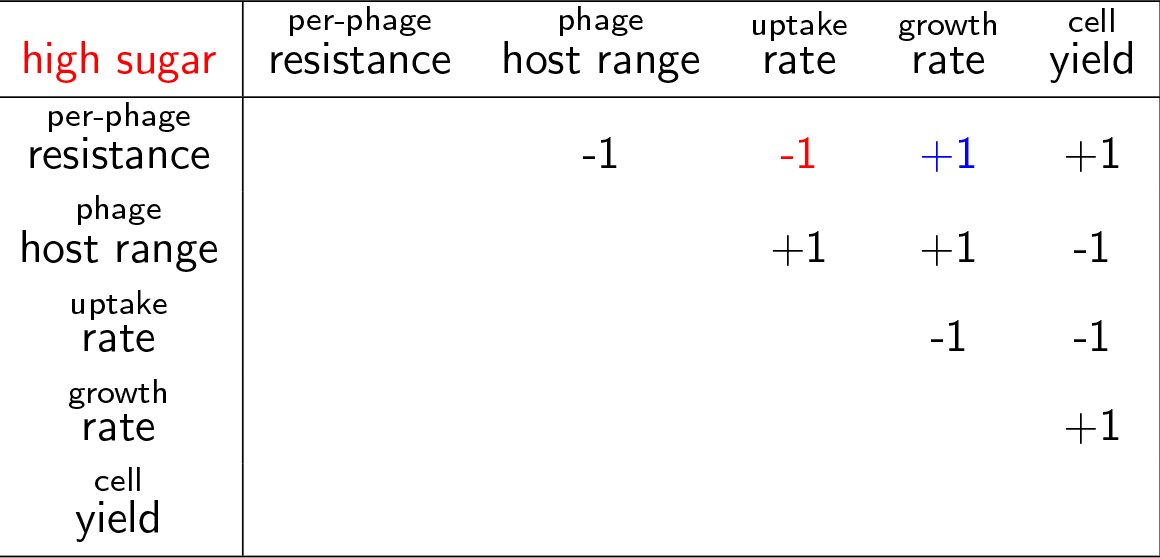
Parity relationships showing tradeoffs (−1) and tradeups (+1) between the 5 traits studied in the main text. This table is not universal as some of these relationships are environment-dependent, but this is a possible interaction at sufficiently high maltotriose supply.

| high sugar | per-phage<br>resistance | phage<br>host range | uptake<br>rate | growth<br>rate | cell<br>yield |
| --- | --- | --- | --- | --- | --- |
| per-phage<br>resistance |  | -1 | -1 | +1 | +1 |
| phage<br>host range |  |  | +1 | +1 | -1 |
| uptake<br>rate |  |  |  | -1 | -1 |
| growth<br>rate |  |  |  |  | +1 |
| cell<br>yield |  |  |  |  |  |

**Supplementary Table S5:** Parity relationships showing tradeoffs (−1) and tradeups (+1) between the 5 traits studied in the main text. This table is not universal either but what might be expected at low maltotriose supply.

| low sugar | per-phage<br>resistance | phage<br>host range | uptake<br>rate | growth<br>rate | cell<br>yield |
| --- | --- | --- | --- | --- | --- |
| per-phage<br>resistance |  | -1 | -1 | -1 | +1 |
| phage<br>host range |  |  | +1 | +1 | -1 |
| uptake<br>rate |  |  |  | +1 | -1 |
| growth<br>rate |  |  |  |  | -1 |
| cell<br>yield |  |  |  |  |  |

## Footnotes

1 As the algorithm proceeds the superscript *K* represents the well-matched points we want to keep, whereas *L* represents the poorly matched points we want to to lose.

2 This condition says that this particular point-matching attempt has been seen before (the algorithm has returned to a prior guess), it will therefore enter a loop if we continue, so instead output the best match found thus far and stop.

